# Deubiquitinating Enzyme Amino Acid Profiling Reveals a Class of Ubiquitin Esterases

**DOI:** 10.1101/2020.05.11.087965

**Authors:** Virginia De Cesare, Daniel Carbajo Lopez, Peter D. Mabbitt, Odetta Antico, Nicola T. Wood, Satpal Virdee

## Abstract

The reversibility of ubiquitination by the action of deubiquitinating enzymes (DUBs) serves as an important regulatory layer within the ubiquitin system. Approximately 100 DUBs are encoded by the human genome and many have been implicated with pathologies including neurodegeneration and cancer. Non-lysine ubiquitination is chemically distinct and its physiological importance is emerging. Here we couple chemically and chemoenzymatically synthesized ubiquitinated lysine and threonine model substrates to a matrix-assisted laser desorption/ionization time-of-flight (MALDI-TOF) mass spectrometry-based DUB assay. Using this platform, we profile two-thirds of known catalytically active DUBs for threonine esterase and lysine isopeptidase activity and find that most DUBs demonstrate dual selectivity. However, with two anomalous exceptions, the ovarian tumor domain (OTU) DUB class demonstrate specific (iso)peptidase activity. Strikingly, we find the Machado Josephin Domain (MJD) class to be unappreciated non-lysine DUBs with highly specific ubiquitin esterase activity that rivals the efficiency of the most active isopeptidases. Esterase activity is dependent on the canonical catalytic triad but proximal hydrophobic residues appear to be general determinants of non-lysine activity. These findings suggest that non-lysine ubiquitination is an integral component of the ubiquitin system, with regulatory sophistication comparable to that of canonical ubiquitination.

## Introduction

Ubiquitination impacts on almost all cellular processes and is carried out by a multienzyme cascade involving E1 activating enzymes (E1), E2 conjugating enzymes (E2) and E3 ligases (E3s)^1,2^. E3s confer substrate specificity and can broadly be classified into two main classes. The largest class consist of RING E3s which use an adapter-like mechanism to facilitate direct transfer of ubiquitin (Ub) from upstream thioester-linked E2 (E2~Ub) to substrate^3^. On the other hand, engagement of E2~Ub by HECT-like E3s results in formation of a thioester-linked E3 intermediate that carries out substrate transfer autonomously^4,5^. Conventionally, Ub is linked to the *ε*-amino group of lysine side chains by an isopeptide bond, or less frequently, it can be appended to the *α*-amino group of proteins via a regular peptide bond^1^. Multiple residues within Ub itself can also become ubiquitinated allowing the formation of Ub polymers with distinct linkage topologies which can mediate different cellular processes^6^. Ubiquitination is a dynamic modification and is reversed by the action of deubiquitinating enzymes (DUBs). Approximately 100 DUBs have been identified in humans and are assigned to 7 distinct classes^7^. For the majority of DUBs, substrate specificity is poorly understood with most biochemical insights gained thus far coming from studies towards isopeptide-linked Ub polymers. Alterations in substrate ubiquitination are often the molecular basis for pathology and DUBs have become attractive therapeutic targets^7^.

Although ubiquitination is typically a lysine-specific posttranslational modification, the RING E3 MIR1 encoded by Kaposi’s sarcoma associated-herpes virus, can evade host immune responses by carrying out ubiquitination of cysteine within major histocompatibility complex class I (MHC I) molecules^8,9^. This promotes their endocytosis and lysosomal degradation. It was subsequently shown that MIR1 from murine g-herpes virus also ubiquitinates MHC1 molecules, but targets serine, threonine and lysine residues and promotes their degradation by endoplasmic reticulum-associated degradation (ERAD)^10^. However, the adapter-like mechanism demonstrated by RING E3s relies upon the active site of the E2 to mediate transfer chemistry. This grants E2s with the important ability to direct ubiquitination to specific sites within a substrate^1,11^. It was subsequently demonstrated that murine MIR1 functions with the poorly studied mammalian E2 UBE2J2, which was shown to possess cellular serine/threonine esterification activity^12^.

In further support of the physiological importance of non-lysine ubiquitination, HECT-like E3s also possess intrinsic esterification activity. MYCBP2/Phr1 has important roles in neural development and programmed axon degeneration^13^, and is has highly selective threonine esterification activity^5^. Furthermore, the E3 HOIL-1 has fundamental roles in immune signalling^14^, and forms ester-linkages with serine/threonine residues within Ub polymers and protein substrates^15^.

The emerging evidence that non-lysine ubiquitination has important roles across a range of fundamental cellular process such as viral infection, ERAD, axon degeneration and immune signalling, places urgent emphasis on establishing which of the ~100 DUBs might confer Ub esterase activity and serve as negative regulators of this chemically distinct form of ubiquitination. A small panel of DUBs have been tested for activity against an ester-linked substrate which indicated that certain DUBs do possess esterase activity and this need not be mutually exclusive with isopeptidase activity^16^. However, comprehensive, DUB profiling, across multiple classes, remains to be carried out.

Here, we synthesize model substrates consisting of threonine/serine that are ester-linked to Ub. Using a high-throughput matrix-assisted laser desorption/ionization time-of-flight (MALDI-TOF) DUB assay^17^, we profile two thirds of known active Ub DUBs for selectivity towards linkage chemistry (lysine isopeptide versus threonine ester). Our findings show that the vast majority of DUBs demonstrate isopeptidase and esterase activity with comparable kinetics. Isopeptidase versus esterase activity is largely inherent to DUB class as USP and UCH DUBs displayed little preference for linkage chemistry. On the other hand, the OTU class were largely dedicated isopeptidases. Two exceptions were TRABID and the virally encoded DUB, vOTU. Strikingly, the MJD class demonstrated pan-selective threonine and serine specific esterase activity. We show that esterase specificity is maintained towards model peptide substrates. We also demonstrate that *in vitro,* the E2 UBE2J2 possesses selective esterification activity, as inferred by its auto modification profile, which is specifically reversed by the MJD member JOSD1. Using chemically synthesized fluorescent substrates we quantify the catalytic efficiency of JOSD1 and find it to be a highly efficient Ub esterase (*k_cat_/K_M_* = 3.5 x 10^4^ M^-1^s^-1^) with a efficiency comparable to that of the most efficient isopeptidases. Taken together, our findings further support the biological significance of non-lysine ubiquitination and demonstrate that its regulatory sophistication is comparable to that of canonical ubiquitination. The complementary activity profiles of certain OTU DUBs with that of JOSD1 might also allow them to be used as research tools for dissecting the emerging prevalence of non-lysine substrate ubiquitination. Our ester-linked model substrates should also facilitate the development of robust assays for inhibitor screening against MJD members.

## Results

### DUB esterase and isopeptidase activity profiling

To determine DUB activity and specificity toward either ester or isopeptide bonds, we employed a previously developed MALDI-TOF DUB assay^17^, and the model substrates Ub-Lysine (Ub-Lys) and Ub-Threonine (Ub-Thr) (**Figure 1**). We reasoned that the differences in the free energy associated with recognition of a ubiquitinated lysine side chain versus a ubiquitinated threonine side chain would be minor and selective esterase activity would manifest through enhanced catalytic turnover (*k_cat_*), rather than reduced Michaelis constant (*K_M_*). If this were the case, these substrates should informatively report on *k_cat_* and hence approximate DUB selectivity within the context of protein substrates.

**Fig. 1.**
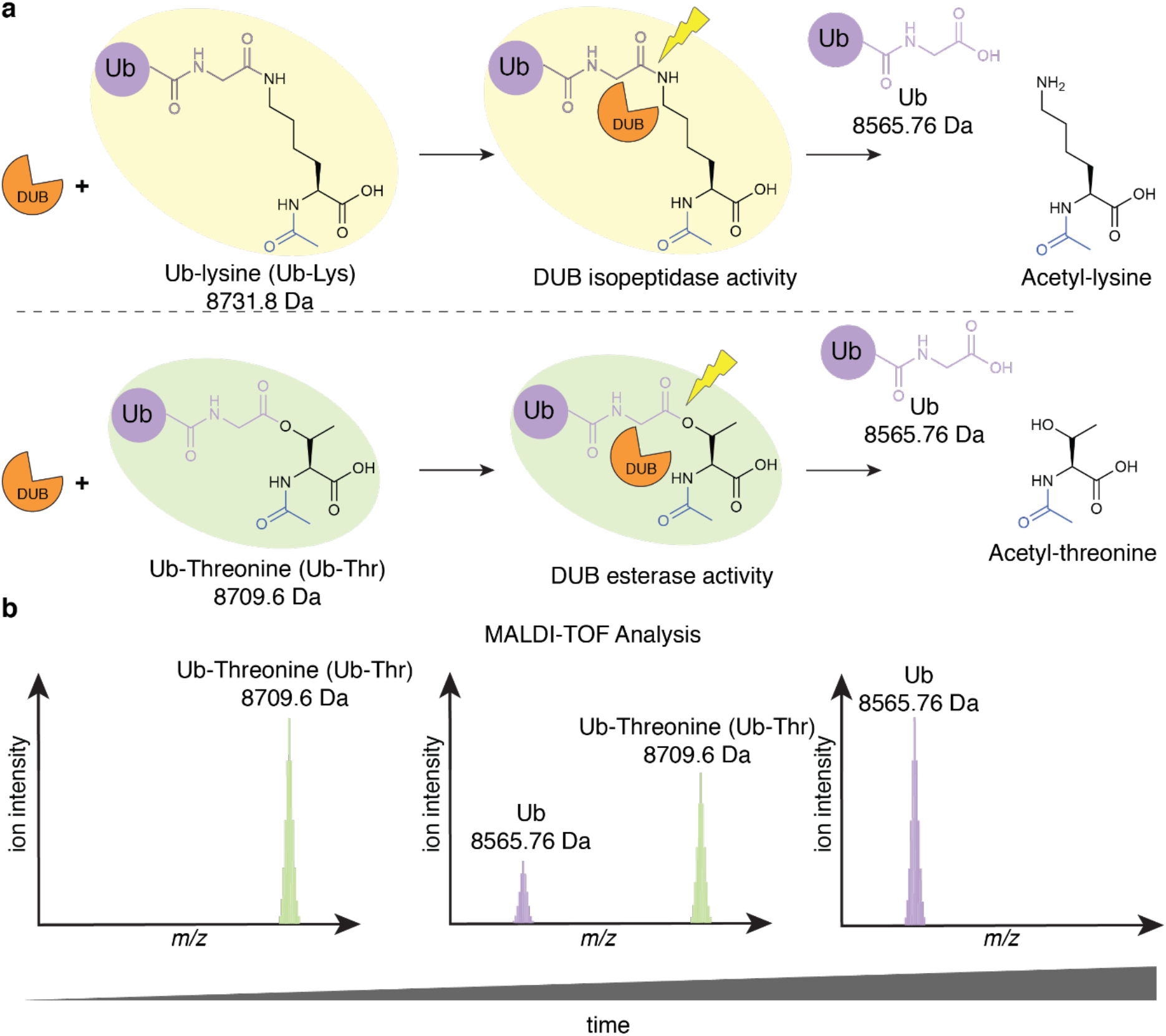
MALDI-TOF-based methodology for profiling DUB isopeptidase/esterase activity. DUBs are incubated either with ubiquitin-lysine (Ub-Lys) or ubiquitin-threonine (Ub-Thr). Reactions are quenched by addition of trifluoracetic acid (2 %), spotted on an AnchorChip 1536 target and analysed by MALDI-TOF mass spectrometry. DUB isopeptidase or esterase activity releases free lysine or threonine and generates native Ub (8565.76 Da) in a time-dependent manner. Ion intensity ratios for peaks corresponding to Ub-Lys (8731.8 Da)/Ub-Thr (8709.6 Da) versus Ub (8565.76 Da) are used for qualitative assessment of % cleavage.

Ub-Lys was chemically prepared using a modified implementation of GOPAL technology^18^ (**Figure S1**). Ub-Thr was chemoenzymatically prepared using a reconstituted E1-E2-E3 cascade based on the RING-Cys-Relay E3 machinery from MYCBP2^5^ (**Figure S2**). For both amino acid substrates, the *α*-amino group was acetylated, which helped mirror the peptide context the model substrates were reflective of, and also prevented potential O-N acyl transfer of Ub-Thr to a peptide-linked species^19^.

We screened a panel of 53 recombinant DUBs belonging to all 7 known DUB families (**Figure 2 a, b**)^7^: ubiquitin-specific protease (USP), ovarian tumor domain (OTU), ubiquitin C-terminal hydrolase (UCH), JAB1/MPN/Mov34 metalloenzyme (JAMM), Machado Josephin Domain (MJD), motif interacting with Ub-containing novel DUB family (MINDY) and zinc finger with UFM1-specific peptidase domain protein (ZUFSP). DUBs were incubated with Ub-Lys and Ub-Thr and as positive controls, they were also incubated with an alternative Ub-derived substrate (either an isopeptide-linked diubiquitin or Ub with a C-terminal peptide-linked adduct) known to be processed by the DUB under investigation (**Fig S3 and Table S1**). However, for the DUBs JOSD1, OTU1, OTUD6A and OTUD6B, a readily accessible substrate which is cleaved remains to be identified. For quantification and normalization purposes, the ratio of the area of the substrate ion intensity signal (Ub-Lys or Ub-Thr) and the area of the product signal (ubiquitin) were recorded and extrapolated to a standard curve, based on defined substrate/product ratios, enabling calculation of % substrate cleavage (**Figure S4**). By comparing the % of cleavage of Ub-Lys versus Ub-Thr as function of time we determined activity and specificity toward the two model substrates **(Figure 2b)**.

**Fig. 2.**
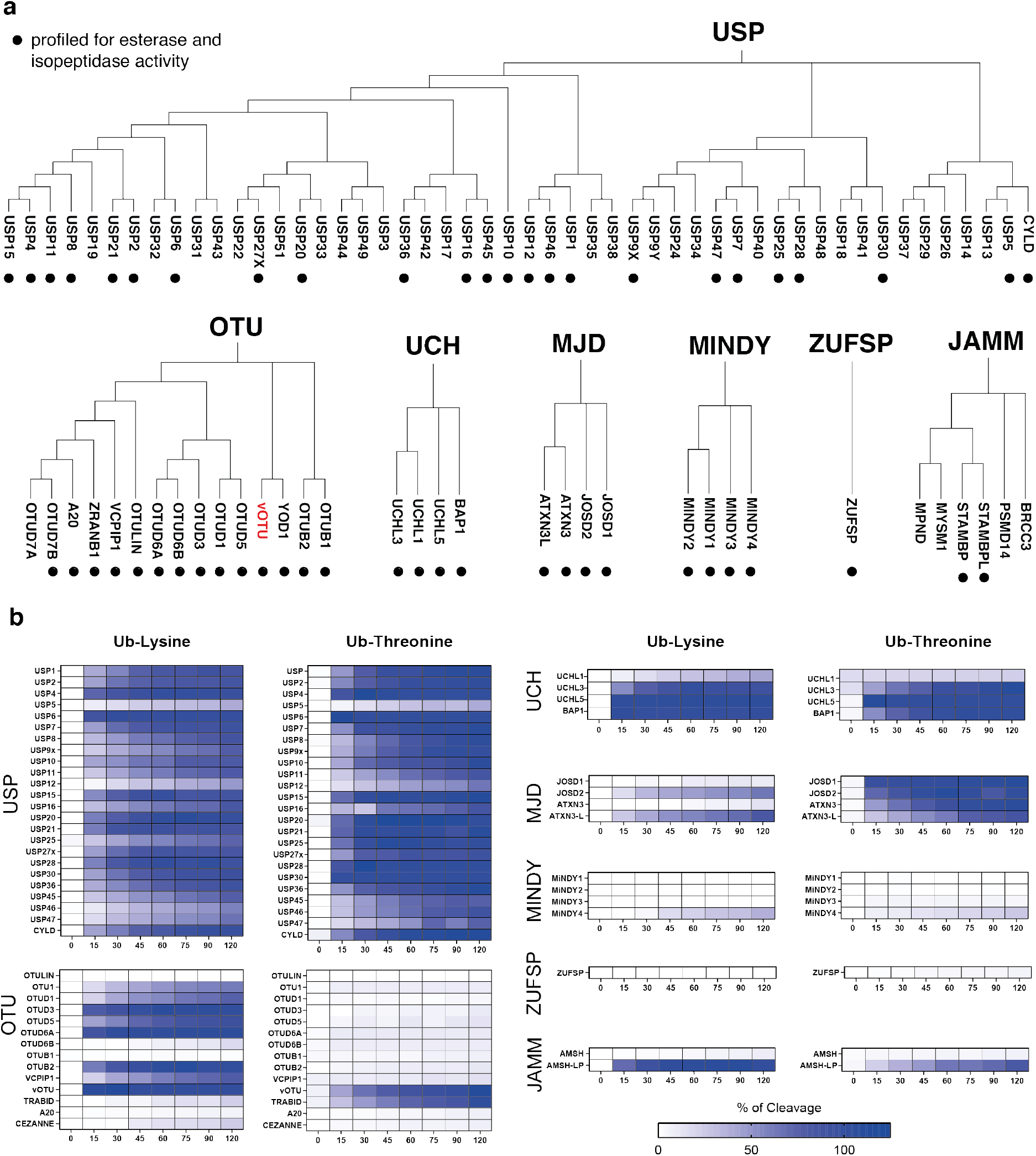
DUB esterase and isopeptidase screen by MALDI-TOF mass spectrometry. **(a)** Phylogenetic classification of deubiquitinating enzymes based on their catalytic domains. Only DUBs that are active and recognise ubiquitin are displayed. vOTU (red) catalytic domain exists within Protein L, which is encoded by Crimean-Congo hemorrhagic fever virus. DUBs annotated with a solid circle correspond to those profiled for activity in this study. **(b)** A panel of 53 DUBs were tested for their activity toward model substrates ubiquitin-lysine and ubiquitin-threonine. Reactions were then quenched by addition of trifluoroacetic acid (2 %) at the relevant time points and spotted on to a 1536 AnchorChip target plate followed by MALDI-TOF analysis^17^. Results are reported as % of cleavage in a scale from white (no activity) to dark blue (100 % substrate consumption). Employed DUB concentrations are specified in Table S1.

### Esterase and isopeptidase selectivity is largely inherent to DUB classification

We find that DUBs belonging to the USP and UCH family in general cleave both Ub-Lys and Ub-Thr substrates with comparable kinetics (**Figure 2b**). The USPs are the largest class consisting of ~60 members whereas UCHs are a smaller classification consisting of only 4 members^7^. The only USPs with negligible activity towards either substrate are USP5 and USP12. However, lack of USP5 activity is anticipated as it is only functional towards polyubiquitin with an intact free Ub C-terminus^20^. In the case of the UCH class, UCHL1 is found to only cleave the isopeptide-linked substrate while UCHL3, UCHL5 and BAP1 show no appreciable preference between the two model substrates (**Figure 2b**).

In contrast to USP and UCH classes, OTU family members invariably demonstrate efficient lysine isopeptidase activity towards our model substrate but negligible threonine esterase activity (**Figure 2b**). Two notable exceptions are vOTU and TRABID. In the context of this assay, TRABID demonstrates selective threonine esterase activity whereas vOTU demonstrates robust activity towards both substrates (**Figure 2b**). The DUB vOTU is encoded by the deadly human pathogen, Crimean Congo haemorrhagic fever virus. In addition to its ability to hydrolyze four out of six tested isopeptide-linked ubiquitin polymer types^21^, it has also been shown have relaxed substrate scope as it also removes the ubiquitin-like modifier ISG15^21^. Thus, the observation that vOTU demonstrates high isopeptidase and threonine esterase activity implies that relaxation of its substrate scope also extends to Ub linkage chemistry and that non-lysine ubiquitination may promote mammalian anti-viral responses more broadly.

TRABID has been implicated with Wnt and immune signalling^22,23^, and has efficient isopeptidase activity in the context of Lys29- and Lys33 linked Ub polymers^18,24^. However, in our assay TRABID has negligible isopeptidase activity towards Ub-Lys, consistent with that observed for other, albeit peptide-linked, small molecule substrates^25^. Thus, our observation that TRABID has high activity towards Ub-Thr implies that its esterase activity is more promiscuous than its isopeptidase activity and that a significant proportion of its physiological substrates may in fact be non-lysine ubiquitination sites.

The Machado-Josephin Domain family is a small class of DUBs consisting of four members^26^. Unlike the other DUB classifications which are found in all eukaryotes, MJD DUBs are absent in yeast, perhaps due to a specific demand of higher eukaryotes. Strikingly, with the exception of ATXN3-L all MJD DUBs demonstrate preferential threonine esterase activity (**Figure 2b**). This is particularly notable for JOSD1 where isopeptidase activity is negligible but quantitative cleavage of the Ub-Thr substrate is observed after the first time point. Similarly, Josephin-2 (JOSD2) cleaves both substrates but has a significant preference for Ub-Thr substrate over the lysine counterpart. Ataxin-3 also cleaves Ub-Thr more efficiently than Ub-Lys (**Figure 2b**).

We also tested the recently discovered MINDY and ZUFSP classes of DUB. The MINDY class consists of 4 members, which demonstrate *exo* (cleaving from the distal end) activity towards extended Lys48 linked Ub polymers^27^. ZUFSP consists of a single founding member (ZUFSP/ZUP1) and is specific for Lys63-linked Ub polymers and is involved in DNA repair^28–31^. Consistent with activity of these DUBs being dependent on polyUb linkage context, negligible activity is observed towards either of our model isopeptide or ester-linked substrates (**Figure 2b**).

Unlike the other DUB classes identified thus far which are cysteine isopeptidases/peptidases, the JAMM class of DUBs are metalloproteases^26^. Two of the six functional JAMM class DUBs included in our panel are AMSH and AMSH-LP, which both have been shown to have specific activity towards isopeptide-linked Lys63 Ub polymers, with comparable efficiency. Interestingly, under the enzyme concentrations employed, AMSH displays no detectable esterase nor isopeptidase activity towards the model substrates whereas AMSH-LP is active against both model substrates (**Figure 2b**)^32^.

### Validation of selective USP and OTU isopeptidase activity

To validate the activity profiles determined by the MALDI-TOF assay format we initially prepared a fluorescent model substrate where the *α*-amino group of threonine is labelled with TAMRA (Ub-Thr-TAMRA) (**Figure 3a, b**). The fluorescent amino acid was linked to Ub via an ester bond using the chemoenzymatic strategy adopted earlier. For comparison we used commercially available isopeptide-linked Ub-Lys-TAMRA-Gly (**Figure 3c**). Esterase or isopeptidase activity, respectively, would cleave the fluorescent amino acid from Ub allowing continuous and quantitative measurement of DUB activity by fluorescence polarization (FP)^33^. For orthogonal testing in the FP assay we selected the USP class DUB USP2 and vOTU, both of which demonstrate high isopeptidase and esterase activity (**Figure 2b**). We also selected OTUB2, OTUD3 and OTUD6A, which demonstrate isopeptidase specificity, and TRABID, which demonstrates selective threonine esterase activity despite reports of it being an efficient isopeptidase (**Figure 2b**). Under these alternative assay conditions USP2 maintains dual specificity with comparable observed rates for isopeptidase and threonine esterase activity (0.18 min^-1^ and 0.13 min^-1^, respectively) (**Figure 3d)**. The DUB vOTU also maintains dual specificity albeit with a preference for Ub-Lys-TAMRA-Gly (**Figure 3e**). Again, consistent with the MALDI-TOF assay, the OTU DUBs OTUD3, OTUB2, OTUD6A exhibit isopeptidase specificity as no threonine esterase activity is detected (**Figure 3f-h**). However, despite demonstrating threonine esterase activity towards the small acetylated threonine substrate, TRABID is only weakly active against Ub-Thr-TAMRA and, unexpectedly, similarly so towards the Ub-Lys-TAMRA-Gly (**Figure 3i**). To ascertain if the C-terminal glycine residue in the isopeptide linked substrate might be affecting the ability to process the isopeptide-linked substrate, relative to the less hindered Ub-Thr-TAMRA substrate, we also prepared an analogous Ub-Lys-TAMRA substrate (**Figure S5**), and found activity to be comparable towards both (**Figure S6**). This is indicative of the TAMRA fluorophore present in Ub-Thr-TAMRA interfering with recognition by TRABID.

**Fig. 3.**
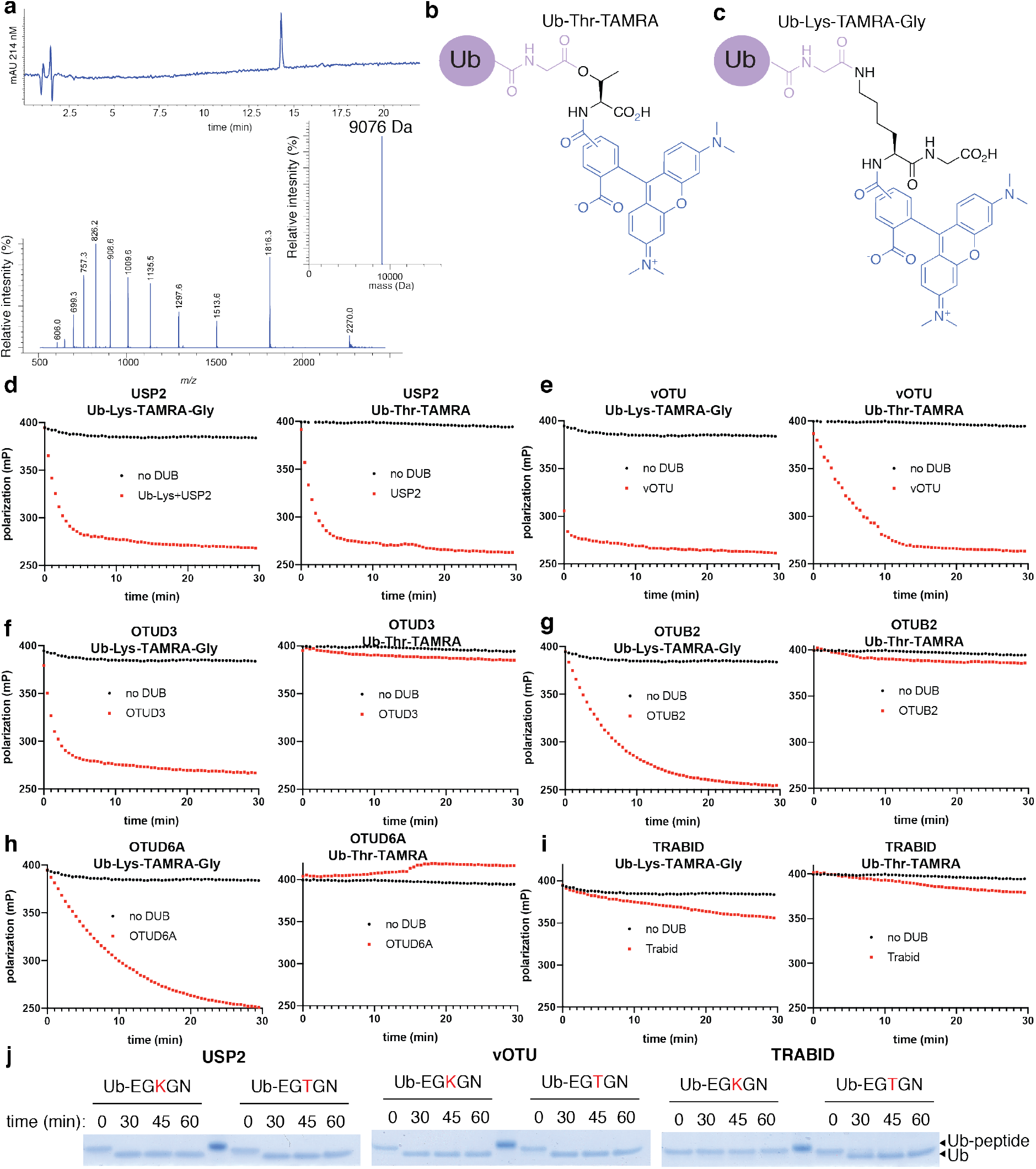
Validation of selected DUB activity towards Ub-Lys-TAMRA-Gly and Ub-Thr-TAMRA by continuous FP-based assay, and ubiquitinated model lysine and threonine peptides by gel-shift assay. **a)** LC-MS characterization data for Ub-Thr-TAMRA: HPLC chromatogram for purified Ub-Thr-TAMRA monitoring at 214 nm, electrospray ionization mass spectrum for Ub-Thr-TAMRA and deconvoluted mass spectrum for Ub-Thr-TAMRA (theoretical mass = 9078.65; observed mass = 9076 Da). **b)** Chemical structure of the fluorescent Ub-Thr-TAMRA structure **c)** Chemical structure of commercial Ub-Lys-TAMRA-Gly. **d)** Consistent with the MALDI-TOF data, the USP2 (0.125 μM) demonstrates comparable lysine isopeptidase and threonine esterase activity. **e)** The virally encoded DUB, vOTU (1.5 μM), has both lysine isopeptidase and threonine esterase activity with an apparent selectivity for the isopeptide-linked model substrate. **f-h)** OTUD3, OTUB2 and OTUD6A (1.5 μM) demonstrate selective lysine isopeptidase activity. **i)** Inconsistent with the MALDI-TOF data using the simple model substrates Ub-Lys and Ub-Thr, TRABID (1.5 μM) demonstrates both weak isopeptidase and esterase activity. **j)** To ascertain whether the observed DUB activity profiles were observed in a peptide context, activity was assessed towards unlabelled ubiquitinated model peptides (Ub-EGKGN and Ub-EGTGN). Consistent with both MALDI-TOF data using unlabelled amino acid substrates and FP data using TAMRA labelled substrates, both USP2 and vOTU (0.75 μM) demonstrate robust isopeptidase and esterase activity. Consistent with the MALDI-TOF data towards unlabelled Ub-Lys and Ub-Thr, TRABID (0.75 μM) demonstrates negligible lysine isopeptidase activity but robust threonine esterase activity.

### TRABID demonstrates selective threonine esterase activity towards a peptide substrate

The unexpected esterase activity towards Ub-Thr, and the inconsistent results obtained from this substrate and Ub-Thr-TAMRA, prompted us to prepare alternative substrates based on model peptides (Ac-EGXGN-NH2 where X = K or T) that were either isopeptide-linked or ester-linked to ubiquitin. DUB activity towards these peptide substrates would be more reflective of a native protein substrate and could be assessed by electrophoretic shift upon SDS-PAGE analysis. Ubiquitinated peptides were prepared using a reconstituted E1-E2-E3 cascade based on a constitutively active RING E3 (RNF4)^34^, or MYCBP2, and were purified by reversed phase - high performance liquid chromatography (RP-HPLC) (**Figure S7–8**). We initially tested USP2 and vOTU. Consistent with the activity profiles towards the Ub-Lys and Ub-Thr substrates, USP2 and vOTU cleaves both the isopeptide-linked and threonine-linked peptide substrates within our first time point, supporting the likelihood that the bispecific isopeptidase and esterase activity demonstrated by the vast majority of USP DUBs and vOTU would extend to protein substrates (**Figure 3j**).

Consistent with the initial MALDI-TOF assay data using non-fluorophore labelled amino acid substrates, TRABID demonstrates threonine esterase activity within the peptide context but no lysine isopeptidase activity (**Figure 3j**). This suggests that whilst TRABID has efficient isopeptidase activity towards distinct polyUb linkages^18^, it reinforces our earlier assertion that its esterase activity is indeed more promiscuous and TRABID is likely to have unappreciated substrates that are ester-linked to Ub.

### Machado-Josephin Domain Family (MJD) DUBs have pan-selective ubiquitin esterase activity

The Machado-Josephin family (also referred to as Josephins) are a small class of DUBs consisting of four members (ATXN-3, ATXN3-L, JOSD1 and JOSD2). The founding and most-studied member is Ataxin-3 (ATXN3)^35,36^. This protein is encoded by the gene responsible for the neurological condition spinocerebellar ataxia type-3, or Machado-Joseph Disease, from which the MJD class takes its name. Machado-Joseph Disease is an autosomal dominant neurodegenerative disorder caused by polyQ tract expansion^37^. All MJD proteins share a common globular catalytic cysteine protease domain of ~180 amino acids, known as the Josephin domain. Ataxin-3 and Ataxin3-L sequences consist of the Josephin domain and a disordered C-terminal tail where the polyQ tract and two or three ubiquitin-interacting motifs (UIMs) are located. The latter bind polyUb chains (**Figure 4a**)^36,38^. JOSD1 and JOSD2, on the other hand, consist of little more than the catalytic Josephin domain.

**Fig. 4.**
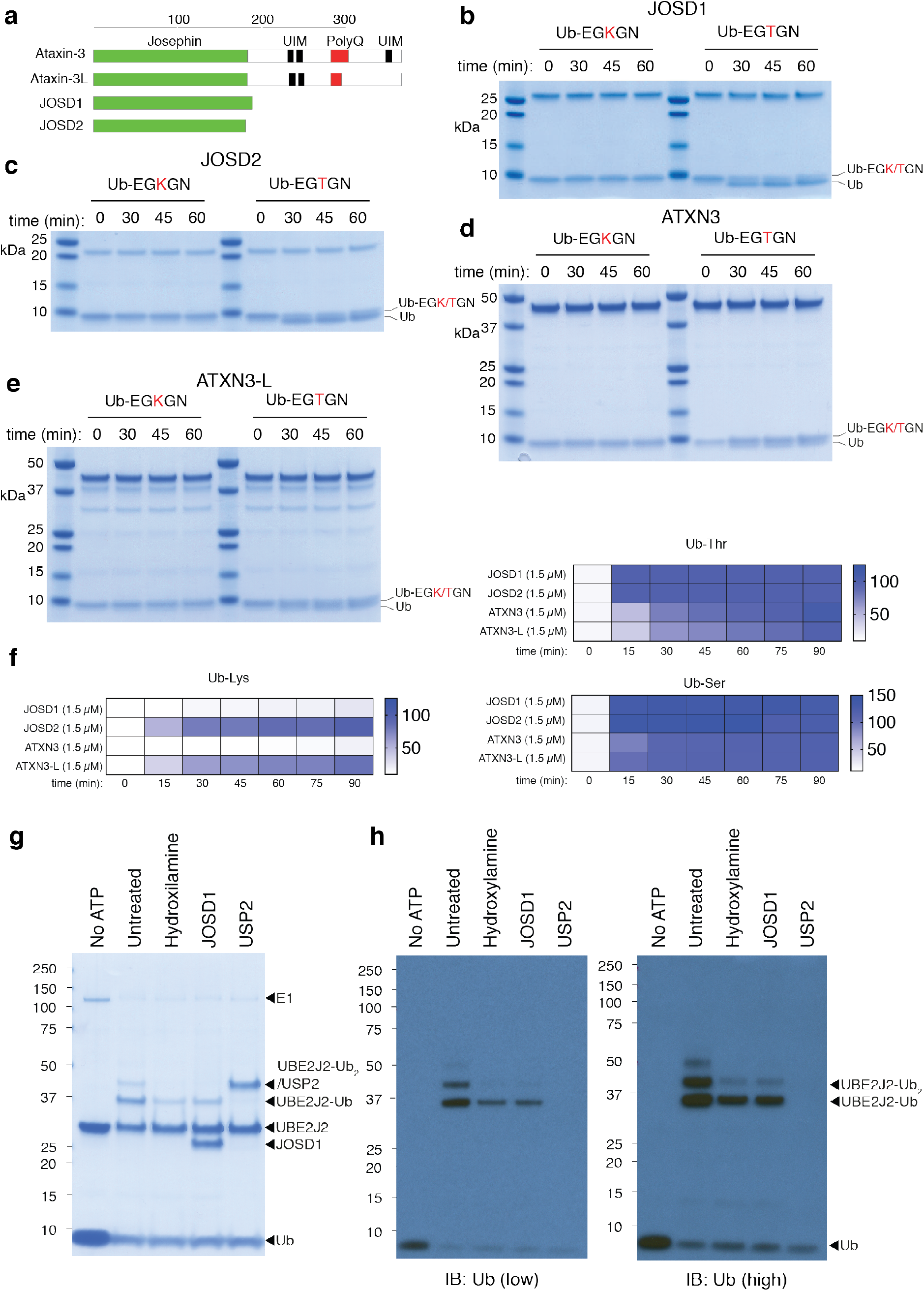
Comprehensive assessment of MJD DUB selective esterase activity towards peptides and the non-lysine auto-modification product of the E2 conjugating enzyme UBE2J2. **a)** Domain architecture of Machodo Josephin DUBs. **b)** Assessment of JOSD1 selective threonine esterase activity towards Ub-EGKGN and Ub-EGTGN model peptide substrates. **c**) Assessment of JOSD2 selective threonine esterase activity towards model peptide substrates. **d**) Assessment of ATXN3 selective threonine esterase activity towards model peptide substrates. **e**) Assessment of ATXN3-L selective threonine esterase activity towards model peptide substrates. **f**) MALDI-TOF assay data demonstrating that with the exception of ATXN3-L, all MJD family members demonstrate pan selective threonine/serine esterase activity towards model Ub-Lys, Ub-Thr and Ub-Ser substrates. **g)** Enzymatic reconstitution in phosphate buffered saline (pH 7.5) with UBE2J2 (20 μM), E1 (200 nM), Ub (50 μM) and ATP (2 mM). Reaction was incubated for 30 min at 37 °C and E2 loading was terminated by addition of Compound 1 E1 inhibitor (25 μM)^47^. Samples were then treated with buffer control or DUB (3 μM) for 30 min at 30 °C. Reactions were terminated with SDS loading buffer containing 2-mercaptoethanol (BME) or BME plus hydroxylamine (0.5 M) and incubated for 30 min at 37 °C prior to SDS-PAGE analysis. **h)** High and low exposures for anti-Ub immunoblot of samples analysed in **g**.

JOSD1 is implicated with membrane dynamics^39^, cancer chemoresistance^40^ and antiviral response^41^, yet its physiological substrates are poorly defined^40,41^. It has been reported that a fraction of JOSD1 exists in a mono-ubiquitinated form which preferentially localizes to the plasma membrane^39^. Importantly, it has recently been shown that elevated JOSD1 levels are found in gynaecological cancers and this correlates with poor prognosis^40^. JOSD1 also promotes chemoresistance by stabilizing anti-apoptotic myeloid cell leukaemia 1 (MCL-1). We too observe ubiquitin modified JOSD1 when transiently overexpressed in HEK293 cells (**Figure S9**) which has been shown to have a modest activating effect on its ability to cleave polyUb species^39^. We could also immunologically detect JOSD1 in various mouse tissue with highest levels of expression observed in the heart, liver, kidney and spleen in mouse tissue lysates (**Figure S9**). Although MJD DUBs have been shown to have isopeptidase activity towards polyubiquitin substrates, this has not been quantified and the qualitative data that does exist involves high micromolar concentrations of DUB and/or lengthy incubation times (e.g. 16-20 hours)^39^. To reconcile the sluggish isopeptidase kinetics it has been proposed that ATXN3 may serve as a cellular timer^42^. Another, possibility is that the precise nature of the physiological substrates is yet to be determined.

Our observation that MJD DUBs have potent threonine esterase activity raised the possibility they were an unappreciated class of specific ubiquitin esterase. To explore this possibility further we tested the four MJD DUBs for activity against the peptide substrates. JOSD1 demonstrates efficient threonine esterase activity but no detectable isopeptidase activity (**Figure 4b, c**). We also find that in the peptide context, ATXN3 and ATXN3-L demonstrate specific threonine esterase activity, albeit weaker than JOSD1 and JOSD2 (**Figure 4d, e**). These findings imply that all 4 MJD DUBs have highly selective, if not specific, esterase activity towards our degenerate peptide substrates. Unlike JOSD1/2, ATXN3 and ATXN3L possess UIM domains that recognise polyUb chains^36^. Thus, the attenuated esterase activity of these DUBs towards our monoubiquitinated substrates might be balanced by polyUb binding in cells, allowing them to mediate preferential esterase activity within the context of polymeric Ub chains.

To address whether MJD esterase activity was specific for threonine we chemoenzymatically prepared ubiquitinated serine substrate (Ub-Ser) using the previously employed conjugation system based on MYCBP2 (**Figure S10**). MJD DUBs were then screened in parallel, using the MALDI-TOF assay, for activity against Ub-Lys, Ub-Thr and Ub Ser. These assays revealed that the MJD class demonstrate both serine and threonine esterase activity with comparable efficiency (**Figure 4f**).

We next assessed whether JOSD1 esterase activity and specificity was maintained in a protein context. The lack of tools for studying non-lysine ubiquitination precluded the development of assays based on a physiological protein substrate. However, the E2 conjugating enzyme UBE2J2 has been reported as having esterification activity, and undergoes auto modification in the presence of E1, Ub and ATP^5,12^. We therefore tested if the auto modifications were ester linked, which would allow the Ub-modified E2 to be used as model protein substrate of DUB esterase activity. We find that a predominant ester-linked Ub adduct is formed, together with a minor (iso)peptide-linked adduct (**Figure 4g, h**). Strikingly, when employed as a model substrate for JOSD1, specific and quantitative esterase activity is observed (**Figure 4g, h**). Furthermore, treatment with USP2 removes both major and minor bands thereby validating our earlier studies showing that USP2 has both esterase and isopeptidase activity (**Figure 4g, h**)

### Quantification of JOSD1 threonine esterase activity

We next benchmarked JOSD1 threonine esterase activity against USP2 using the FP assay with the fluorescent Ub-Lys-TAMRA-Gly and Ub-Thr-TAMRA substrates. USP2 is a highly active DUB (*k_cat_/K_M_* = ~2.5 x 10^5^ M^-1^s^-1^)^43^ and can be used as a research tool to selectively remove Ub from protein substrates by presumed selective isopeptidase activity^44^. Consistent with mass spectrometry and gel-based assays, USP2 readily cleaves Ub from Ub-Lys-TAMRA-Gly and Ub-Thr-TAMRA with similar kinetics (**Figure 5a, b**). However, JOSD1 readily cleaved the Ub-Thr-TAMRA substrate with an observed rate constant comparable to that of USP2 (**Figure 5b**) but exhibited no detectable isopeptidase activity towards Ub-Lys-TAMRA-Gly (**Figure 5a**). To determine if the additional glycine residue in Ub-Lys-TAMRA-Gly was affecting isopeptidase activity we tested all MJD DUBs against our synthesised Ub-Lys-TAMRA substrate and observed equivalent cleavage kinetics (**Figure S6**).

**Fig. 5.**
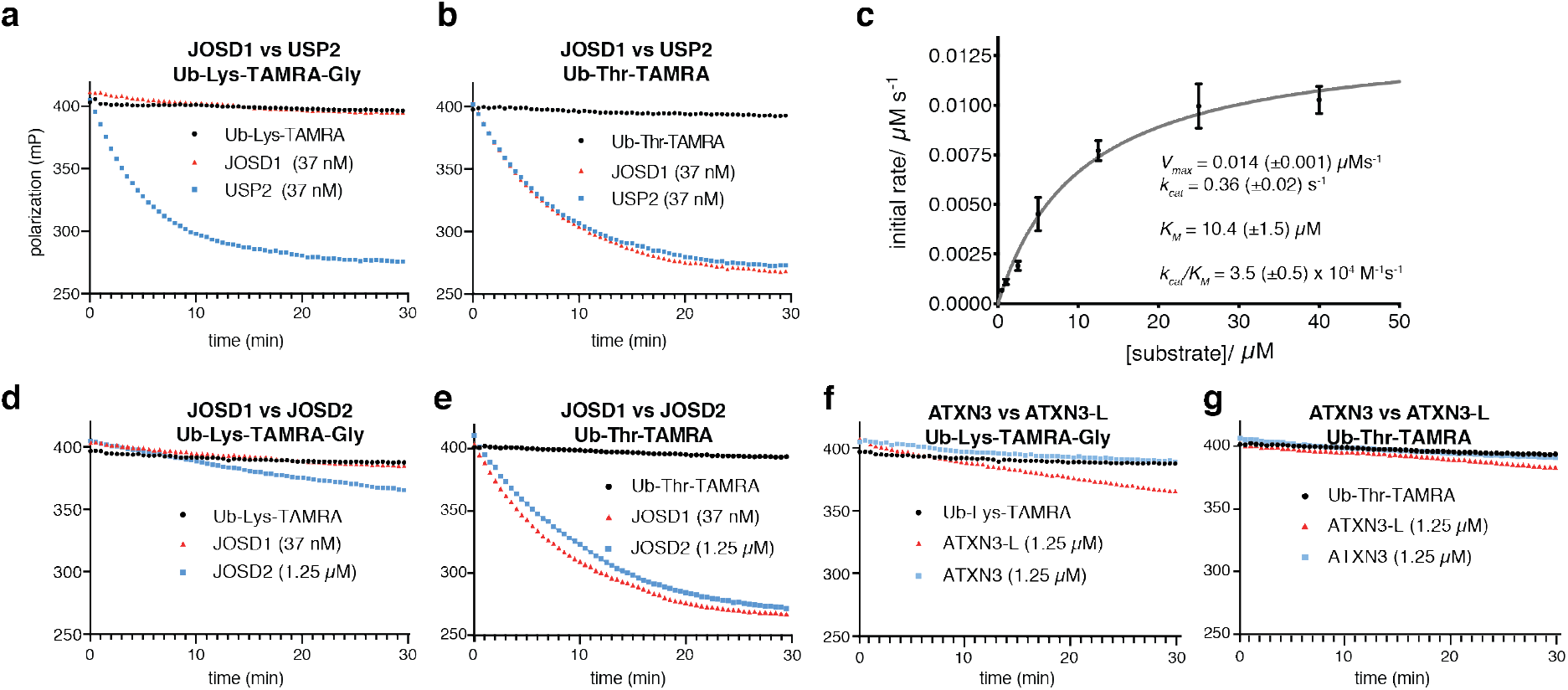
Characterization of Machado Josephin esterase activity by FP-based assay towards model amino acid substrates. **a)** Ub-Lys-TAMRA-Gly/Ub-Thr-TAMRA tested at 250 nM. The USP class DUB USP2 demonstrates efficient isopeptidase activity towards Ub-Lys-TAMRA whereas JOSD1 isopeptidase activity is undetectable. **b**) JOSD1 demonstrates efficient esterase activity towards Ub-Thr-TAMRA with comparable kinetics to that of USP2. **c**) Steadystate Michaelis-Menten analysis for JOSD1 esterase activity towards Ub-Thr-TAMRA (JOSD1 assay concentration was 37 nM). **d**) Negligible JOSD2 isopeptidase activity is also observed towards Ub-Lys-TAMRA (note JOSD2 concentration is 1.25 μM). **e**) JOSD2 is a less efficient esterase than JOSD1 towards Ub-Thr-TAMRA, but cleavage kinetics are comparable when JOSD2 concentration is increased ~30-fold relative to JOSD1. **f**) ATXN3 esterase activity is negligible towards the synthetic Ub-Lys-TAMRA substrate whereas ATXN3-L exhibits weak isopeptidase activity. **g**) Both ATXN3 and ATXN-3L exhibit negligible esterase activity towards Ub-Thr-TAMRA.

To quantify the catalytic efficiency of JOSD1 esterase activity, and determine its catalytic parameters, we carried out Michaelis Menten analysis towards Ub-Thr-TAMRA (**Figure 5c**). Catalytic turnover, *k_cat_*, is 0.36 (±0.02) s^-1^ and Michaelis constant (*K_M_*) is 10.4 (±1.5) μM, indicative of the observed esterase activity being largely driven by *k_cat_*. The resultant specificity constant (*k_cat_/K_M_*) = 3.5 x10^4^ M^-1^s^-1^ is comparable to that of USP21 for the K48 ubiquitin dimer^45^, which is at the high end of the spectrum of kinetically quantified DUB isopeptidase activity. This is consistent with JOSD1 having the potential to mediate dynamic cellular deubiquitination via its esterase activity.

### JOSD1 esterase activity is mediated by the canonical catalytic site

The crystal structure of ATXN3, which has been solved in complex with a ubiquitin suicide substrate probe^42^, allowed us to build a high confidence homology model (99 %) for JOSD1 with Phyre2^46^. In the experimental structure the active site residues are well resolved and although a substrate is absent, residues that would be in immediate proximity of the modified substrate amino acid can be inferred (**Figure 6a, b**). In light of the unexpected esterase activity we first determined whether it was dependent on the established catalytic cysteine within JOSD1. (**Figure 6c**). Canonical JOSD1 activity has been shown to be dependent on C36^39^. We found that mutation a C36A mutant of JOSD1 abolishes activity towards Ub-Thr-TAMRA. Mutation of the histidine belonging to the catalytic triad (H139) also abolishes activity (**Figure 6c**). These results indicate that the catalytic triad centering on C36 in JOSD1, and presumably the homologous residue in other MJD class DUBs, mediates its efficient esterase activity.

**Fig. 6.**
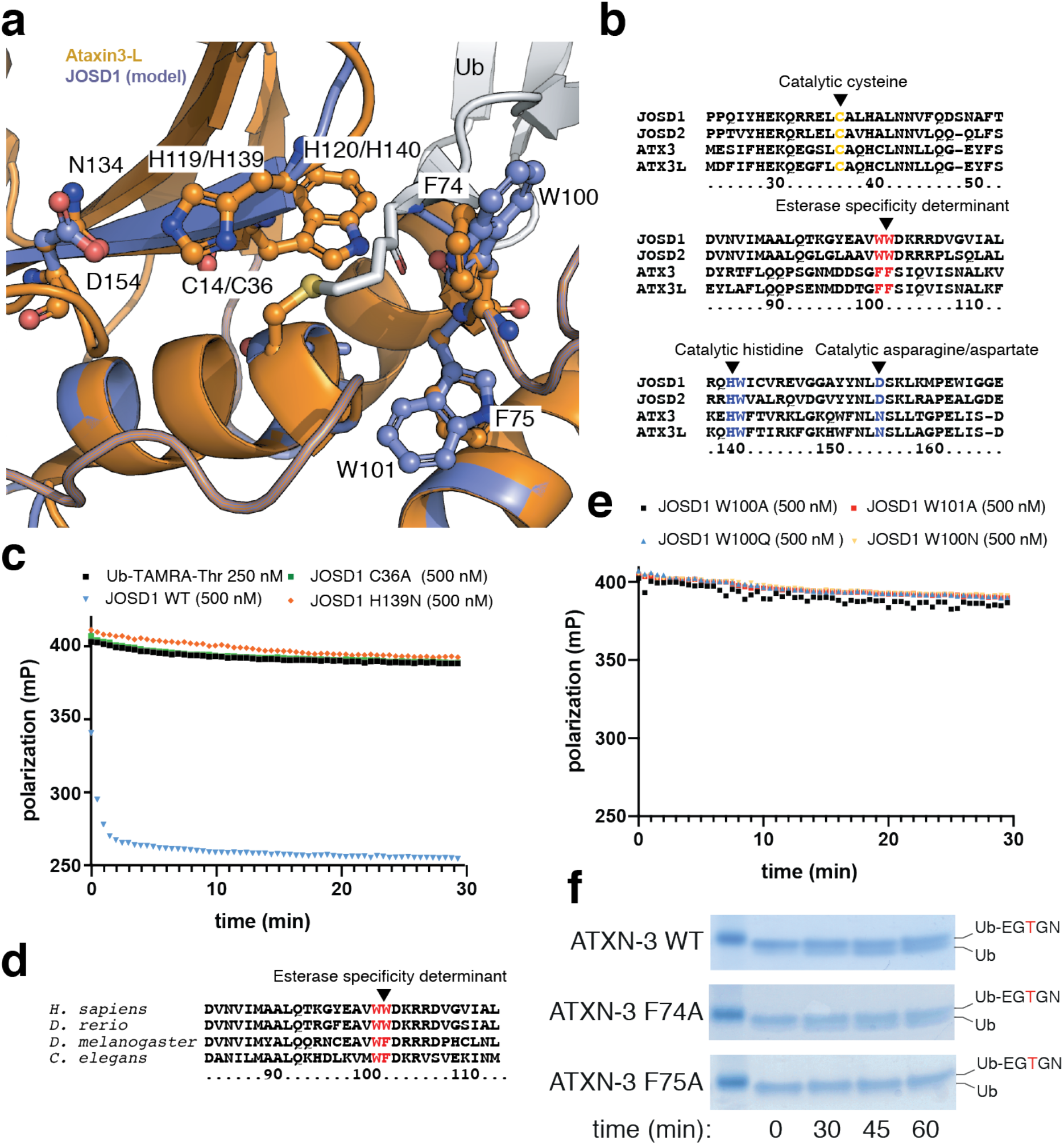
Determinants of MJD DUB esterase selectivity. **a**) Superposition of the active site for a JOSD1 homology model with that of ATXN3-L: Ub complex (PDB ID: 3O65). **b**) Sequence alignment for JOSD1 and ATXN3-L. Catalytic residues and those proposed to be important for esterase activity are highlighted. **c**) JOSD1 esterase activity is dependent on canonical catalytic residues. **d)** Essential W100 and W101 residues are largely conserved but some orthologues have a phenylalanine at position 102 (as for ATXN3 and ATXN3L which have a phenylalanine at both positions). **e)** Hydrophobic Trp residues (W100 and W101) in the JOSD1 active site are required for esterase activity. **f)** Homologous Phe residue F74, but not F75, is required for ATXN3 esterase activity towards a threonine ubiquitinated peptide.

### Assessment of structural basis for MJD DUB esterase activity

To determine catalytic residues that might facilitate the selective threonine esterase activity we studied the experimental ATXN3-Ub structure and its superposition with our JOSD1 homology model **(Figure 6a)**. A conspicuous feature of the ATXN3 and JOSD1 catalytic sites is their hydrophobic nature. The JOSD1 model places two tryptophan residues in the active site whereas ATXN3 has two phenylalanine residues at the equivalent positions (**Figure 6a, b**). In some JOSD1 orthologues the residue equivalent to W101 is a phenylalanine, suggesting that phenylalanine residue can serve a similar function to the tryptophan (**Figure 6d**). We have shown that MYCBP2 E3 ligase threonine esterification activity is dependent on a hydrophobic pocket composed of three phenylalanine residues^5^. Here, the methyl group of the threonine side chain docks into this pocket, which appears to immobilize and activate the threonine side chain to chemistry. However, MYCBP2 also has serine esterification activity, but no detectable lysine aminolysis activity suggesting that the hydrophobic residues in MYCBP2 may contribute to esterification activity more broadly. To test if the hydrophobic residues were contributing to JOSD1 esterase activity we mutated the tryptophan residues and tested activity. Strikingly, both W100A and W101A mutations abolish JOSD1 esterase activity (**Figure 6e**). A W100Q and a W100N mutation also abolish activity implying that hydrophobic character, rather than steric properties, are important (**Figure 6e**). To test if the homologous phenylalanine residues in ATXN3 might have similar function we assayed alanine mutants in the peptide-based gel shift assay. Whilst an ATXN3 F74A mutant has no discernible effect on activity, an F75A mutant abolishes ATXN-3 esterase activity, underscoring the functional relevance of these residues across the MJD class (**Figure 6f**).

## Discussion

Non-lysine ubiquitination is emerging as physiologically important posttranslational modification. As systems-wide technologies for identifying ubiquitination sites are tailored for lysine ubiquitination, the scale of cellular non-lysine ubiquitination remains to be determined. We tested the activity of 53 DUBs against both lysine and non-lysine ubiquitinated model substrates making the current work the most extensive and crossvalidated study on non-lysine DUB activity to date. The results show that isopeptidase versus esterase activity is largely dependent on DUB phylogeny. We found that on the whole, USP and UCH class DUBs mediate both isopeptidase and esterase activity with comparable kinetics. On the other hand, the OTU DUB classification is largely devoid of esterase activity. Amongst the OTU members we studied, TRABID and virally encoded vOTU were exceptions. Towards our model substrates, vOTU demonstrated robust esterase and isopeptidase activity whereas TRABID only conferred esterase activity. However, as TRABID has been shown to mediate efficient isopeptidase activity towards Ub polymers, it would appear that it has the potential to confer dual chemical substrate specificity within cells. Most insights towards DUB substrates have been in the context of polyubiquitin linkage preference. In comparison, knowledge of physiological protein substrates is limited and our findings imply that non-lysine ubiquitination sites must be considered when studying the majority of DUBs (USP, UCH and MJD classes).

The discovery that MJD DUBs, and JOSD1 in particular, having pan-selective Ub esterase activity was striking, suggestive of these DUBs being dedicated to specifically editing non – lysine ubiquitinated substrates. This is particularly evident from our observation that JOSD1 maintains specific esterase DUB activity in the context of the protein substrate UBE2J2. Notably, the molecular basis for UBE2J2 esterification activity, and whether any of the other ~30 E2s might confer esterification activity, remain poorly understood. The modular nature of the RING E3 catalytic mechanism presents the possibility that, in principal, any of the ~600 RING E3s could direct esterification of hydroxy amino acids within substrates. This is in further support of the prevalence of non-lysine ubiquitination being underestimated.

We identified features, within the MJD active site, that are crucial for their activity. The presence of a hydrophobic residues adjacent to the catalytic cysteine appear to serve important roles in mediating selective esterification and esterase activity. The precise role these residues play might be further validated by structural studies. If such hydrophobic residues present within the active site are a universal feature of ubiquitin cascade enzymes (E2s, E3s) that might also have non-lysine activity, then where structural data exists, this feature could potentially be used to predict the existence of such enzymes.

Within the MJD family, ATXN-3 represents the most well studied enzyme as directly related to the development of spinocerebellar ataxia type-3, and its established esterase activity might be of patho-physiological relevance. Within the MJD family, JOSD1 and JOSD2 represent the least studied members. Despite their sequence homology, JOSD1 and JOSD2 seem to have distinct physiological functions and different catalytic efficiencies. JOSD2 has been found able to cleave ubiquitin chains *in vitro* condition while JOSD1 was consistently reported as having low activity or being inactive^39^. These results are consistent with our finding that JOSD2 shows some degree of activity against the isopeptide-linked model-substrate. However, it should be noted that MJD DUB activity towards isopeptide-linked substrates has not been quantified.

The association of JOSD1 with the development of chemoresistance in gynaecological cancer make it subject of a potential biomarker and therapeutic target. However, developing a robust assay for screening JOSD1 inhibitors has been challenging due to the low signal window obtained with conventional (iso)peptide-linked substrates. The substrates produced in our study, notably Ub-Thr-TAMRA, should facilitate the development of revolutionary assay platforms for screening inhibitor screening of MJD DUBs; an emerging class of therapeutic targets. Although only modest activity towards our substrate was observed with ATXN3 and ATXN3-L, the production of an ester-linked polyUb variant may be a more efficient substrate owing to the ubiquitin interacting motifs (UIMs) specific to these members. These could engage the polyUb chain and enhance catalytic efficiency through avidity effects (reduction of *K_M_*). Furthermore, the strikingly specificity and high catalytic efficiency of JOSD1 threonine esterase activity might allow it to be used in combination with DUBs with selective isopeptidase activity (e.g. OTUD3, OTUB2, OTUD6) for diagnosis of Ub linkage chemistry within cellular substrates.

## Acknowledgments

We thank Axel Knebel, Richard Ewan, Clare Johnson and Daniel Fountaine from the MRC Protein Production and Assay Development team, and MRC Reagents and Services, who all contributed to the generation of protein reagents required for the MALDI-TOF DUB assay platform. We are grateful to Ronald Hay for provision of the plasmid encoding the constitutively active RNF4 E3 ligase. This work was funded by UK Medical Research Council (MC_UU_12016/8). We also acknowledge pharmaceutical companies supporting the Division of Signal Transduction Therapy (Boehringer-Ingelheim, GlaxoSmithKline and Merck KGaA).

## Material and Methods

Ubiquitin monomer, BSA, Tris, DMSO and DTT were purchased from Sigma-Aldrich. Ub-Lys-TAMRA was purchased from Boston Biochem (Boston, MA). MALDI-TOF MS 1536 AnchorChip was purchased from Bruker Daltonics (Bremen, Germany). 2’,6’-dihydroxyacetophenone (2,6-DHAP matrix) was purchased from Tokyo Chemical Industry (Product Number: D1716). JOSD1 monoclonal antibody was purchased from ThermoFisher (MA5-25365).

### Synthesis of Ub-Lys and Ub-Thr model substrates

Ub-Lys was prepared using a modification of GOPAL technology, which was developed for the production of defined isopeptide-linked Ub chains^18^. In brief, an excess of *Nα*-acetyl-L-lysine was dissolved in DMSO together with a chemically protected Ub thioester protein in presence of AgNO_3_ and *N*-hydroxysuccimimide as catalyst. After incubation at 23 °C, protein was precipitated with ice-cold diethylether and air-dried. Chemical protecting groups were removed as previously described^18^, and deprotected protein was isolated by ether precipitation. Dried protein was dissolved in denaturing guanidinium chloride buffer and purified by reversed phase high performance liquid chromatography (RP-HPLC)^17^.

Ub-Thr and Ub-Ser were prepared chemoenzymatically in reaction buffer consisting of Na_2_PO_4_ (40 mM), NaCl (150 mM), MgCl_2_ (5 mM), *Nα*-acetyl-L-threonine (50 mM), E1 Uba1 (500 nM), E2 UBE2D3 (10 μM), GST-MYCBP2cat^5^, Ub (50 μM), ATP (5 mM) and TCEP (0.5 mM) in a volume of 2.33 mL. Reaction was incubated at 37 °C for 1-2 h and Ub-Thr was purified by RP-HPLC using a 20 % to 50% gradient over 60 min (buffer A was 0.1 % TFA in H_2_O and buffer B was 0.1% TFA in acetonitrile).

### Synthesis of ubiquitinated peptide model substrates

EGKGN and EGTGN were purchased from Bio-synthesis and resuspended in water to 250 mM final, the pH adjusted to 7-8 using 0.4N NaOH. Ub~EGKGN and Ub Ub~EGTGN were prepared chemoenzymatically using RNF4 and MYCBP2 respectively. RNF4 (10 μM) or GST-MYCBP2cat^5^ (10 μM) were diluted in reaction buffer consisting of Na_2_PO_4_ (40 mM), NaCl (150 mM), MgCl_2_ (5 mM), *Nα*-acetyl-L-threonine (50 mM), E1 Uba1 (500 nM), E2 UBE2D3 (10 μM), Ub (50 μM), ATP (5 mM) and TCEP (0.5 mM). Reaction was incubated at 37 °C for 1-2 h and Ub-EGKGN and UB-EGTGN were purified by RP-HPLC using a 20 % to 50% gradient over 60 min (buffer A was 0.1 % TFA in H_2_O and buffer B was 0.1% TFA in acetonitrile).

### Synthesis of Ub-Lys-TAMRA and Ub-Thr-TAMRA fluorescent substrates

Initially, threonine and lysine were functionalized at the *Nα* position with 5/6-carboxytetramethylrhodamine (TAMRA). A 10-fold molar excess of amino acid (L-threonine or *Ne-(t*-butyloxycarbonyl-)L-lysine) and DIEA, was mixed with 5/6 TAMRA-OSu (Thermofisher #C1171) in DMSO. After agitation for 24 h, reactions were diluted to 10 % DMSO with H_2_O and products were purified by RP-HPLC and lyophilized, yielding threonine-TAMRA and *Nε*-(*t-* butyloxycarbonyl)-Lys-TAMRA. To remove the *t*-butyloxycarbonyl protecting group from the lysine product, material was dissolved in the minimum volume of dichloromethane, which was subsequently diluted to 60 % with TFA. Solvent was removed with a stream of air and repurified by RP-HPLC, yielding Lys-TAMRA. Fluorescent Ub-Lys-TAMRA and Ub-Thr-TAMRA substrates were prepared as described for Ub-Lys and Ub-Thr but *Nα*-acetyl-L-lysine and L-threonine were substituted for Lys-TAMRA and Thr-TAMRA, respectively.

### Protein Expression and Purification

The expression and production of most of the DUBs and^15^N Ubiquitin as previously described^17^. The additional DUBs were expressed as follows. USP11; USP30 (57-517); USP45; UCH-L3, UCH-L5, MINDY2, MINDY3, MINDY4, ZUFSP, Ataxin-3 and Ataxin-3L (WT and mutants) and ZUFSP were cloned into pGEX-vectors, expressed in BL21 DE3 cells. Expression was induced with 0.05 – 0.25mM IPTG and allowed to proceed for 16h at 16°C, followed by affinity purified on GSH-Agarose using standard protocols. Similarly, UCH-L1, USP47, JOSD1 (WT and mutants) and JOSD2, were cloned into pET-vectors, expressed in BL21 DE3 cells and affinity purified on Nickel-Agarose using standard protocols. MINDY1 was cloned into pMex-vector, expressed in BL21 DE3 cells and affinity purified on Amylose-Agarose using standard protocols. USP12 and USP46 were co-expressed with WDR48 / UAF1 as 6His-tagged fusion proteins in Sf21 cells and purified over Ni-NTA agarose using standard protocols. The UCHL-substrate Ubiquitin-Trp was expressed untagged in BL21 DE3 cells and purified like untagged Ubiquitin. Briefly, expression was induced when cells were at OD600 = 1.0 with 1mM IPTG and left for 3.5h at 37°C. The cells were collected in Milli Q water, resuspended and frozen in liquid nitrogen. After thawing, the suspension was sonicated and clarified by centrifugation (38000 x g for 30min at 4°C.). The lysate was titrated to pH 4.3 with 7% perchloric acid and left overnight. Precipitated proteins were removed by centrifugation and the clarified solution was subjected to cation exchange chromatography on a Source S HR10/10 column using a shallow NaCl gradient. Ubiquitin-Trp eluted at 20mS/cm, equivalent to about 180mM NaCl. The protein was concentrated to 50mg/ml and stabilised with Tris pH 8.0 (50mM f.c.).

### Fluorescent Polarization Assay

Ub-K/T-TAMRA (final concentration 0.250 to 40 uM for kinetic calculation, 0.250 uM for standard assay) were diluted in Fluorescence Polarization (FP) Buffer (50 mM Tris-HCl pH 7.5, 150 mM NaCl, 1 mM DTT, 0.01% BSA) aliquoted into a 384-well plate (Corning, black low volume, round bottom). DUBs were diluted at the indicated final concentration in FP Buffer and added into each well. Fluorescence polarization decay was measured with a cycle time of 30s for 60 cycles at 30 °C using plate reader (Pherarastar, BMG Labtech). Normalized parallel and perpendicular fluorescence intensities were used for further calculations. Graphpad Prism 8.0 was used to fit the data into Michaelis-Menten equations and calculate kinetic parameters.

### SDS-page based ubiquitinated peptide cleavage

DUBs (750 nM final concentration) and Ub-EGKGN/Ub-EGTGN (5 uM final concentration) were diluted in 50 mM Tris-HCl pH 7.0, 50 mM NaCl2 and 1 mM DTT. Reaction was incubated for the indicated time points at 30 degree and stopped by adding 1X final LDS-NuPAGE™ Sample buffer. Samples were run on 1mm, 4-12% Bis-Tris Protein Gels for 45 minutes at 200V and blue Coomassie stained.

### MALDI-TOF DUB Assay

Enzymes and substrates were freshly prepared in the reaction buffer (40 mM Tris-HCl, pH 7.6, 5 mM DTT, 0.005% BSA). DUBs were diluted at the indicated concentrations (table 1), 5.8 ul of Assay buffer and 3 uL of diluted enzymes were aliquoted in a Greiner 384 well plate, flat round bottom, low binding. 1.2 uL of either ubiquitin dimers or Ubiquitin~K /~T were added to the reaction mixture at the final concentration of 1.4 and 2.75 μM respectively. The reaction was incubated at 30 °C and stopped adding 2.5 uL of 10% Trifluoroacetic Acid (TFA) at the indicated time points.^15^N labelled ubiquitin was added as internal standard only to the reaction controls using ubiquitin dimers as substrate.

### Target Spotting and MALDI Mass Spectrometry Analysis

MALDI-target spotting and MS analysis was performed similarly as previously described^48^. Briefly, a Mosquito nanoliter dispenser (TTP Labtech, Hertfordshire, UK) was used to mix 1.2 ul of each reaction with 1.2 ul of 7.6 mg/ml 2,5-dihydroxyacetophenone (DHAP) matrix (prepared in 375 ml ethanol and 125 ml of an aqueous 25 mg/ml diammoniumhydrogen citrate). 200 nL matrix/assay mixture was spotted onto an 1536 AnchorChip Plate. Spotted targets were air dried prior to MALDI TOF MS analysis. All samples were acquired as previously reported^48^ on a Rapiflex MALDI TOF mass spectrometer (Bruker Daltonics, Bremen, Germany) equipped with Compass for flexSeries 2.0, FlexControl and FlexAnalysis software (Version 4.0). Peak intensities were exported as .csv file using FlexAnalysis. An *in house* windows batch script was used to report peak intensities into an excel grid with the same geometry as for the MALDI-target. Ubiquitin over^15^N labelled ubiquitin or ubiquitin~K-~T ratios were considered for further calculations. Ubiquitin ~K and ubiquitin ~T standard curves (Sup Fig. 3) were used to translate normalized ratio into % of cleavage.

### UBE2J2-based DUB Activity Assay

UBE2J2 auto-modification reaction was performed by incubating E1 (200 nM), ubiquitin (50 μM) and recombinant His-UBE2J2 (10 μM)^5^, in reaction Buffer (1X PBS, 20 mM MgCl_2_, 2 mM ATP, 1 mM TCEP) for 30 minutes at 30 °C in a final volume of 100 μL. Compound 1 E1 inhibitor was added at final concentration of 25 μM and the reaction incubated for 15 minutes at 30 °C. The reaction was then sub-aliquoted and treated with either JOSD1 (3 μM), USP2 (3 μM) or 1X PBS buffer for 30 minutes at 30 °C. Reactions were stopped by addition of 4X LDS loading buffer (ThermoFisher Scientific) + BME and supplemented with either water or hydroxylamine (0.5 M). Samples where incubated for 30 minutes at 37 °C and resolved on a 4-12 % Bis-Tris gel at 125 V for 1.4 h. Protein bands were either visualized by Comassie stain or by standard Western blot procedure (anti-ubiquitin antibody BioLegend, Cat Number: 646302, Monoclonal, Mouse – 0.1 μg/mL).

### Tissue Harvesting and Lysates preparation

Brain tissues were rapidly excised and the sub-regions microdissection was performed with an ice cooling plate under stereomicroscopy. Upon isolation, brain subregions and peripheral tissues were collected in a single 1.5 ml microcentrifuge tube and snap-frozen in liquid nitrogen. Tissue samples were stored at −80°C until ready for processing. All tissues were weighed and defrosted on wet ice in fivefold mass excess of freshly prepared, ice-cold lysis buffer containing: 50 mM Tris/HCl pH 7.5, 1 mM EDTA pH 8.0, 1 mM EGTA pH 8.0, 1% Triton X-100, 0.25 M sucrose, 1 mM sodium orthovanadate, 50 mM NaF, 10 mM sodium glycerolphosphate, 10 mM sodium pyrophosphate, 200 mM 2-chloroacetamide, phosphatase inhibitor cocktail 3 (Sigma-Aldrich) and complete protease inhibitor cocktail (Roche). Tissue homogenization was performed using a probe sonicator at 4°C (Branson Instruments), with 10% amplitude and 2 cycles sonication (10 seconds on, 10 seconds off). Crude lysates were incubated at 4°C on wet ice 30 min on ice, before clarification by centrifugation at 20,800 x g in an Eppendorf 5417R centrifuge for 30 min at 4°C. Supernatants were collected and protein concentration was determined using the Bradford kit (Pierce).

## Supplementary Material

### Supplementary Figures

**Figure S1.**
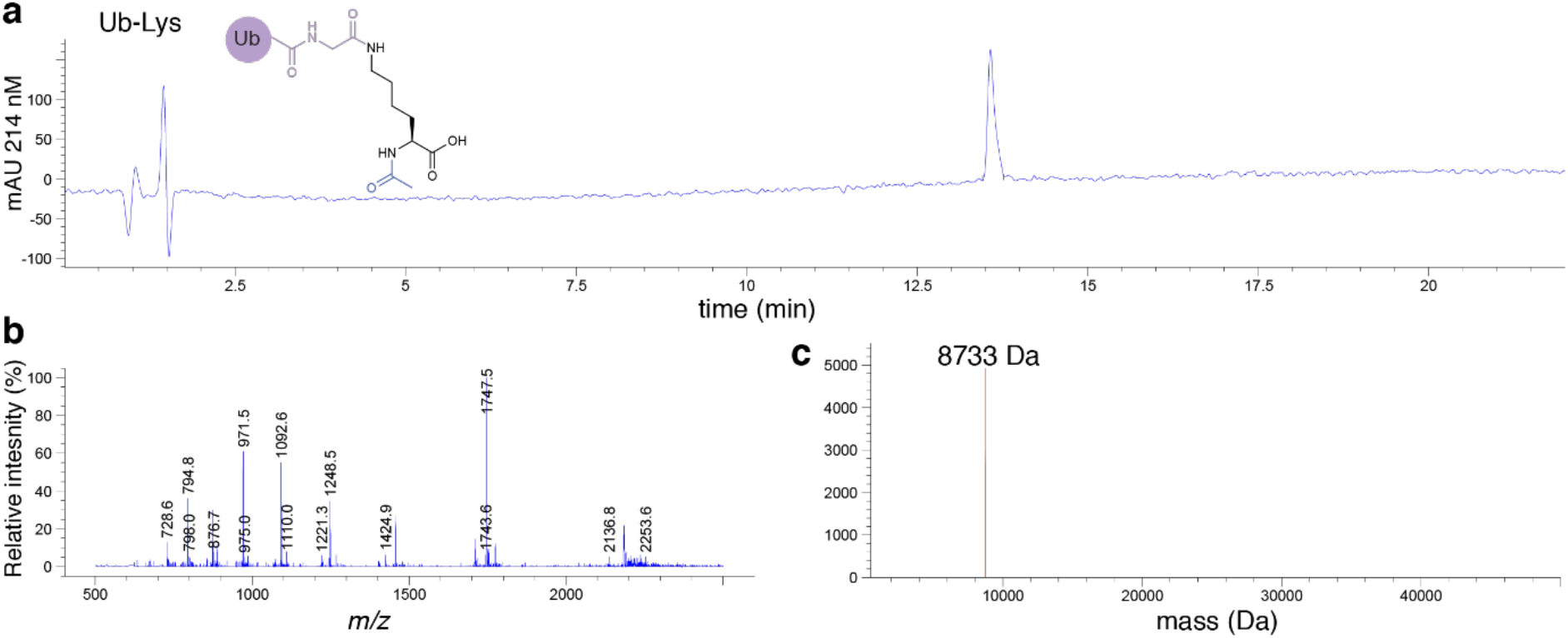
LC-MS characterization of purified Ub-Lys. **(a)** HPLC chromatogram for purified Ub-Lys monitoring at 214 nm. **(b)** Electrospray ionization mass spectrum for Ub-Lys. **(c)** Deconvoluted mass spectrum for Ub-Lys. Theoretical mass = 8735.06; observed mass = 8733 Da.

**Figure S2.**
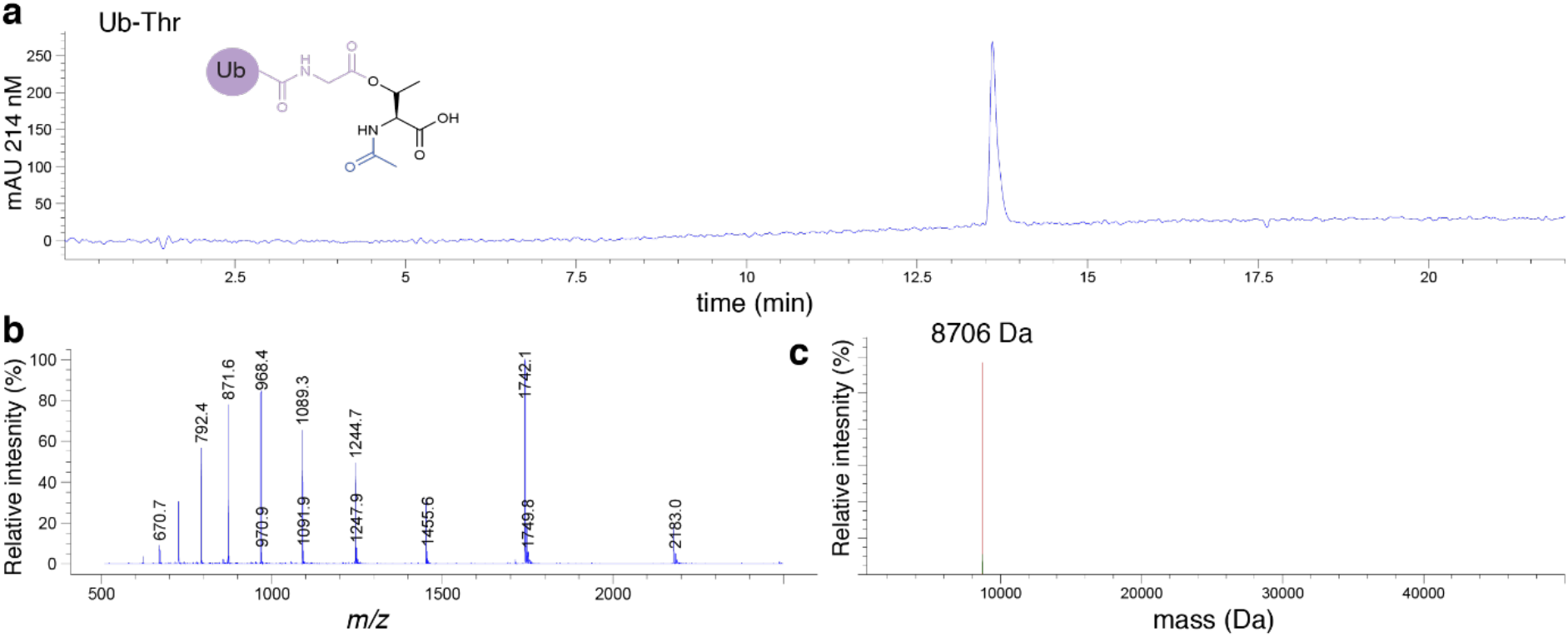
LC-MS characterization of purified Ub-Thr. **(a)** HPLC chromatogram for purified Ub-Thr monitoring at 214 nm. **(b)** Electrospray ionization mass spectrum for Ub-Thr. **(c)** Deconvoluted mass spectrum for Ub-Thr. Theoretical mass = 8708.00.; observed mass = 8706 Da.

**Figure S3.**
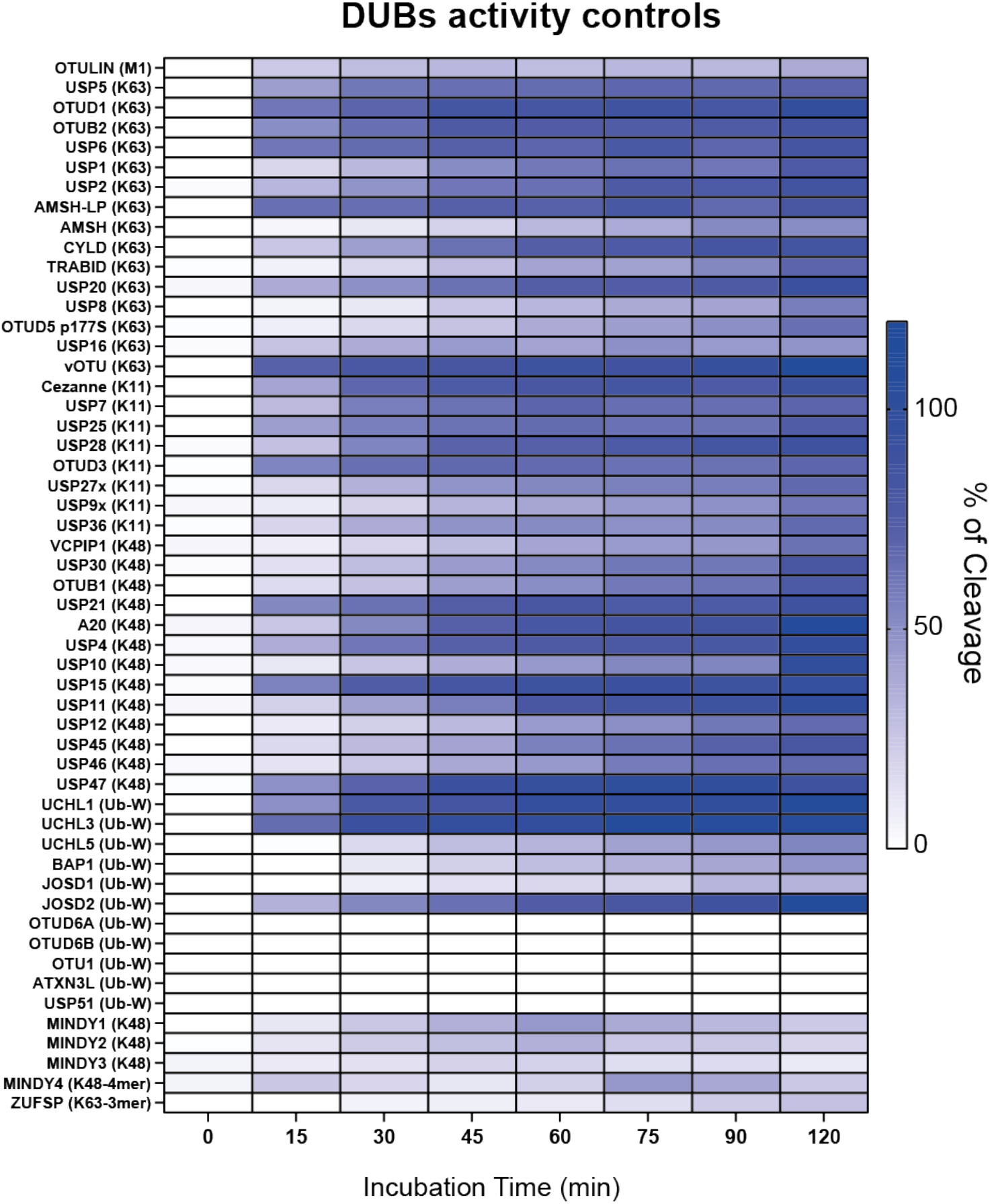
MALDI-TOF DUBs assay: enzymatic activity control: DUBs were tested in parallel for their activity with ubiquitin dimer of linkage type known to be cleaved by the DUB under investigation (indicated as M1-K11-K48-K63). For DUBs that demonstrate negligible activity with ubiquitin dimers Ub with a C-terminal peptide-linked tryptophan was employed. The DUBs JOSD1, OTU1, OTUD6A and OTUD6B demonstrate no detectable activity towards any of the isopeptide-linked Ub dimers nor the peptide-linked substrate.

**Figure S4.**
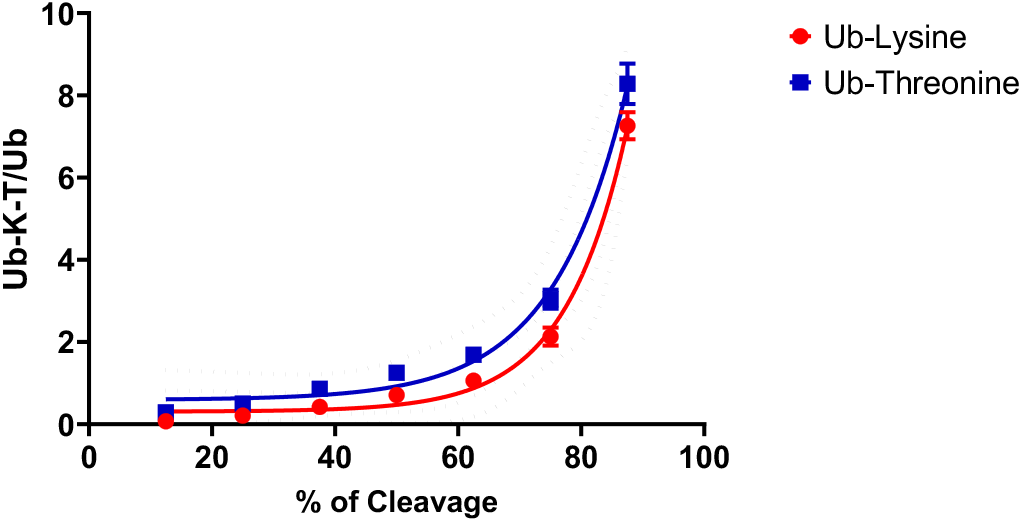
Standard curve for assessment of % substrate cleavage using MALDI-TOF DUB assay. Defined ratios of Ub-Lys (or Ub-Thr) substrate were mixed with free Ub that simulated intermediate states of substrate cleavage. Extrapolation of product/substrate ratios from DUB assays to the standard curve allowed assessment of % substrate cleaved.

**Figure S5.**
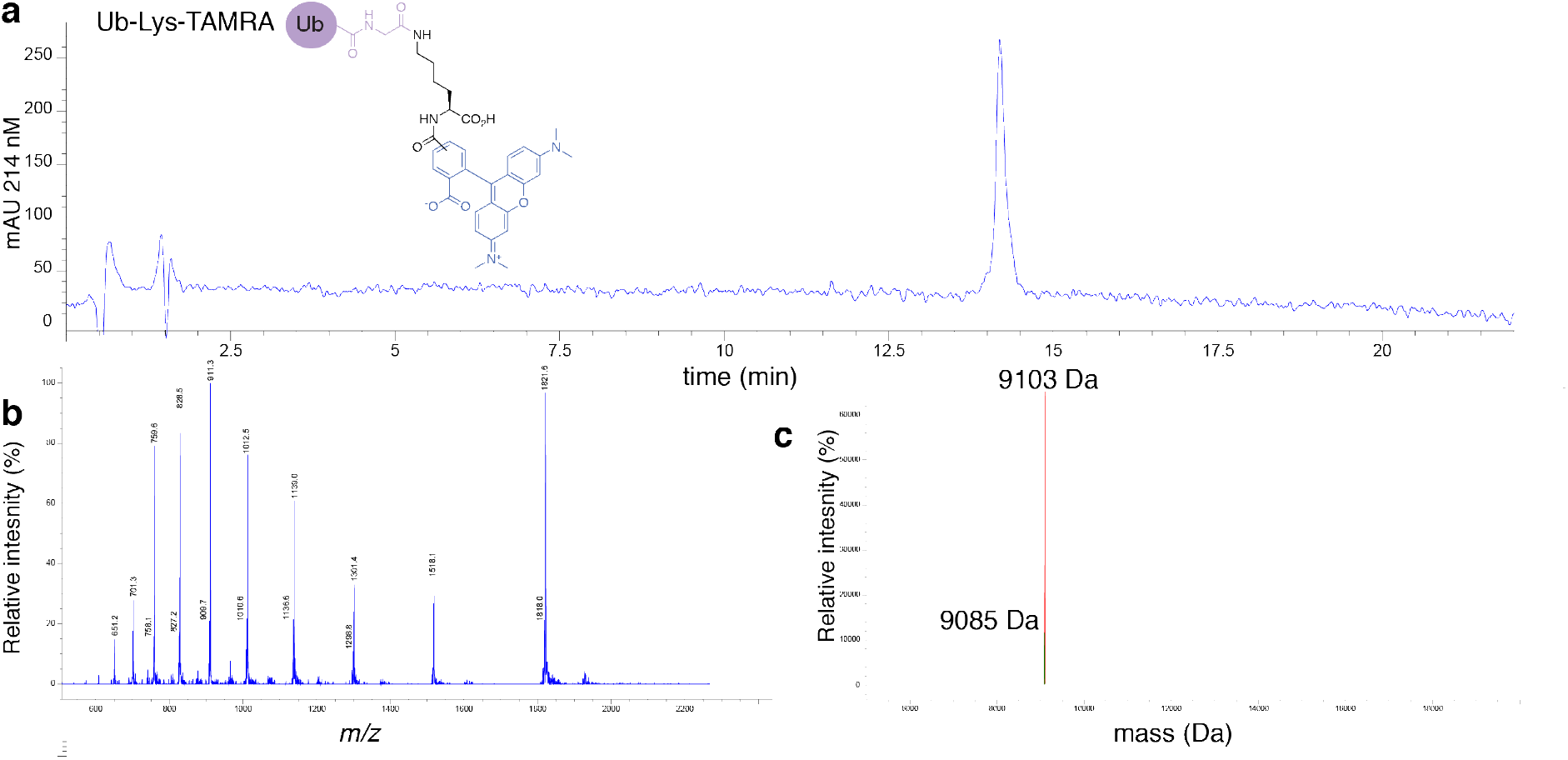
LC-MS characterization of purified Ub-Lys-TAMRA. **(a)** HPLC chromatogram for purified Ub-Lys-TAMRA monitoring at 214 nm. **(b)** Electrospray ionization mass spectrum for Ub-Lys-TAMRA. **(c)** Deconvoluted mass spectrum for Ub-Lys-TAMRA. Theoretical mass = 9105.46; observed mass = 9103 Da. Minor peak with mass of 9085 Da corresponds to a dehydration product.

**Figure S6.**
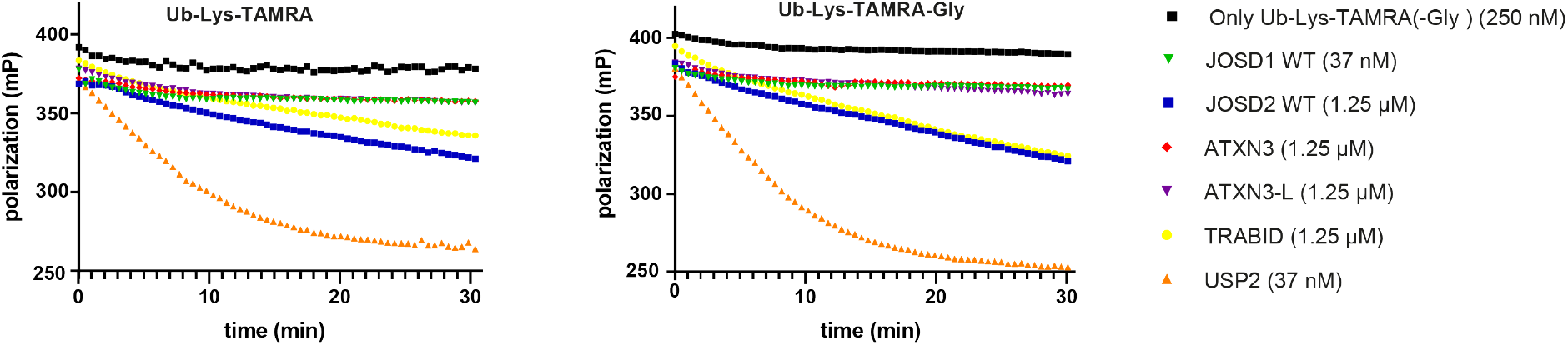
Comparison of activity of selected DUBs towards Ub-Lys-TAMRA and Ub-Lys-TAMRA-Gly.

**Figure S7.**
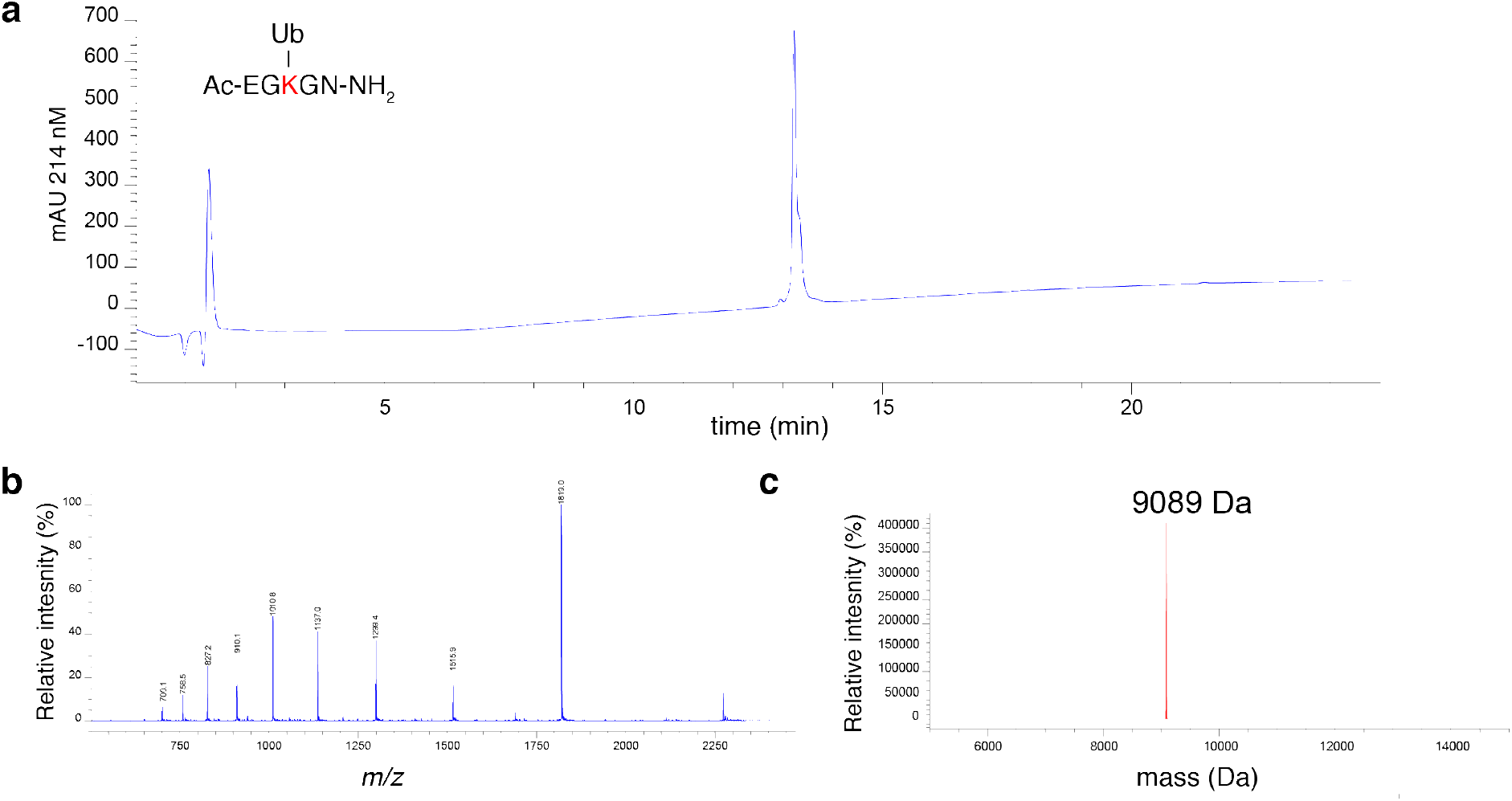
LC-MS characterization of purified Ub-EGKGN. **(a)** HPLC chromatogram for purified Ub-EGKGN monitoring at 214 nm. **(b)** Electrospray ionization mass spectrum for Ub-EGKGN. **(c)** Deconvoluted mass spectrum for Ub-EGKGN. Theoretical mass = 9091.52; observed mass = 9089 Da.

**Figure S8.**
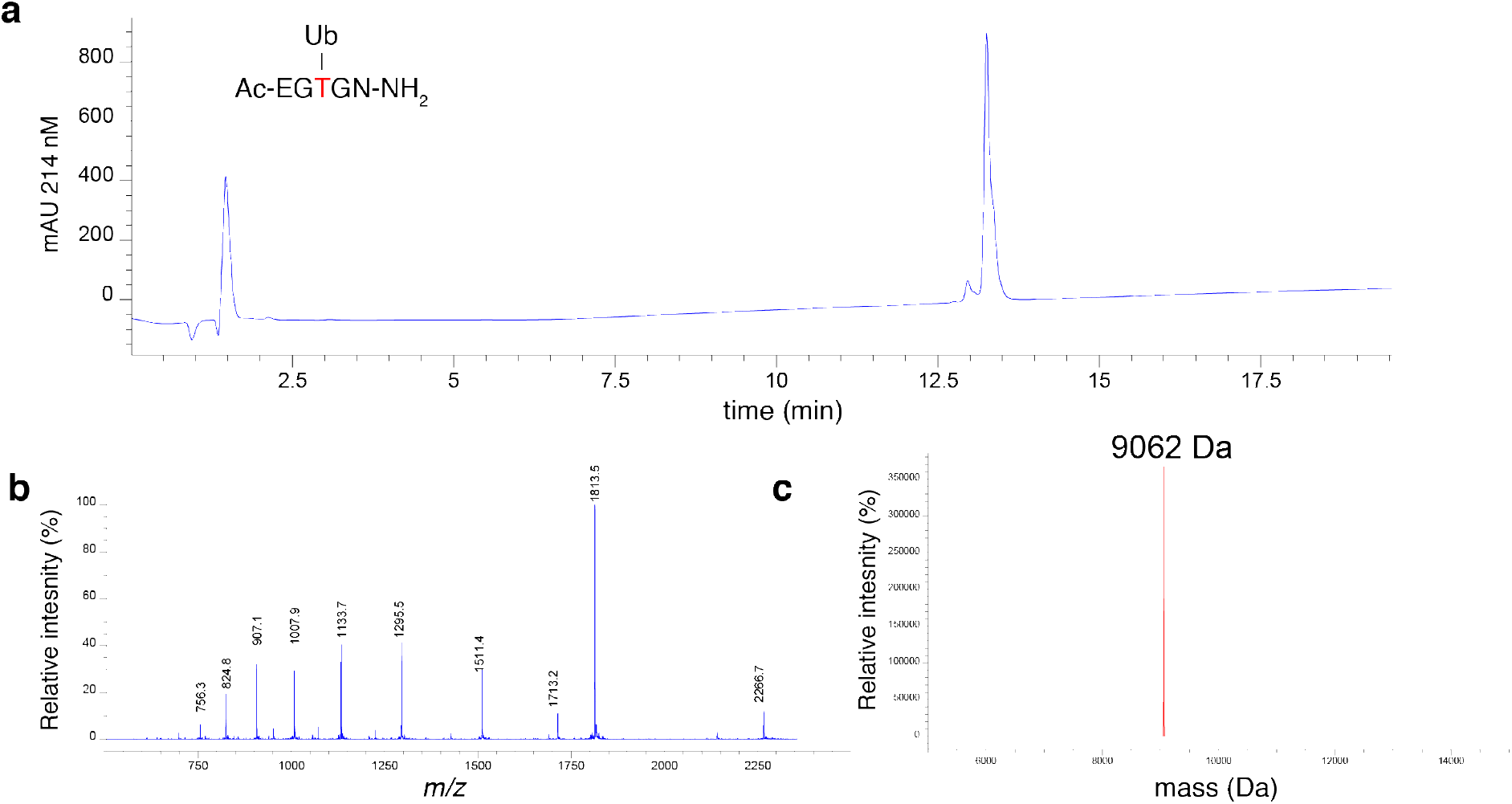
LC-MS characterization of purified Ub-EGTGN. **(a)** HPLC chromatogram for purified Ub-EGTGN monitoring at 214 nm. **(b)** Electrospray ionization mass spectrum for Ub-EGKGN. **(c)** Deconvoluted mass spectrum for Ub-EGTGN. Theoretical mass = 9064.45; observed mass = 9062 Da.

**Figure S9.**
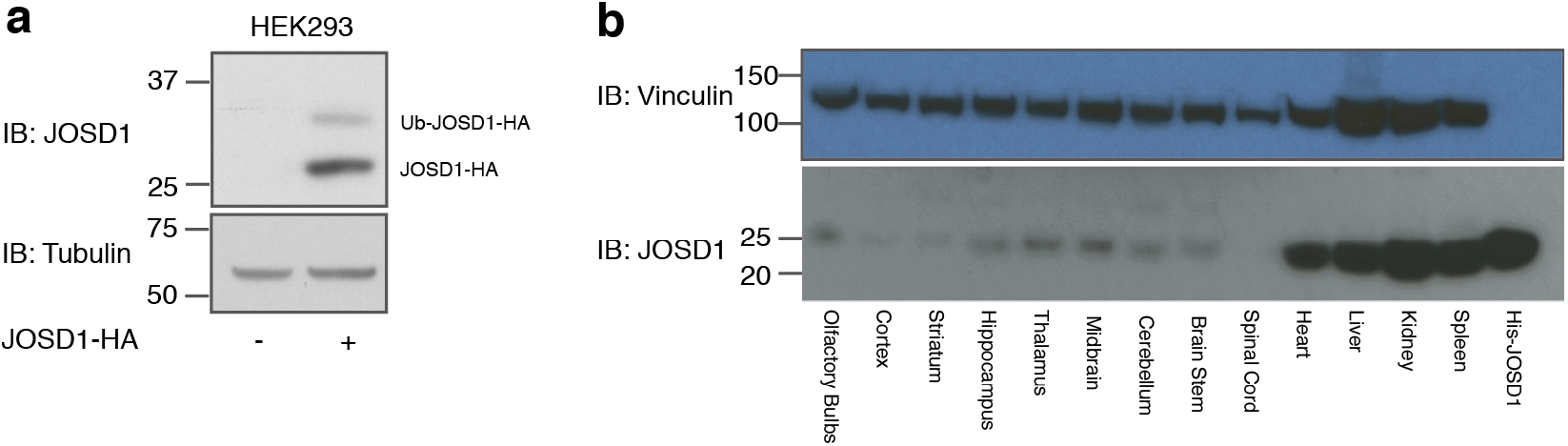
JOSD1 protein expression characterization: **(a)** JOSD1 immunoblot on cultured HEK293 cells. JOSD1 can be detected in HEK293 when transiently expressed. An upper band, corresponding to ubiquitinated JOSD1 is visible. **(b)** Protein lysates from the indicated tissues were immunoblotted with anti-JOSD1 antibody. JOSD1-His recombinantly expressed (30 ng) is used as positive control for accurate detection. JOSD1 is endogenously highly expressed in tissue as hearth, liver, kidney and spleen while less abundant in brain regions.

**Figure S10.**
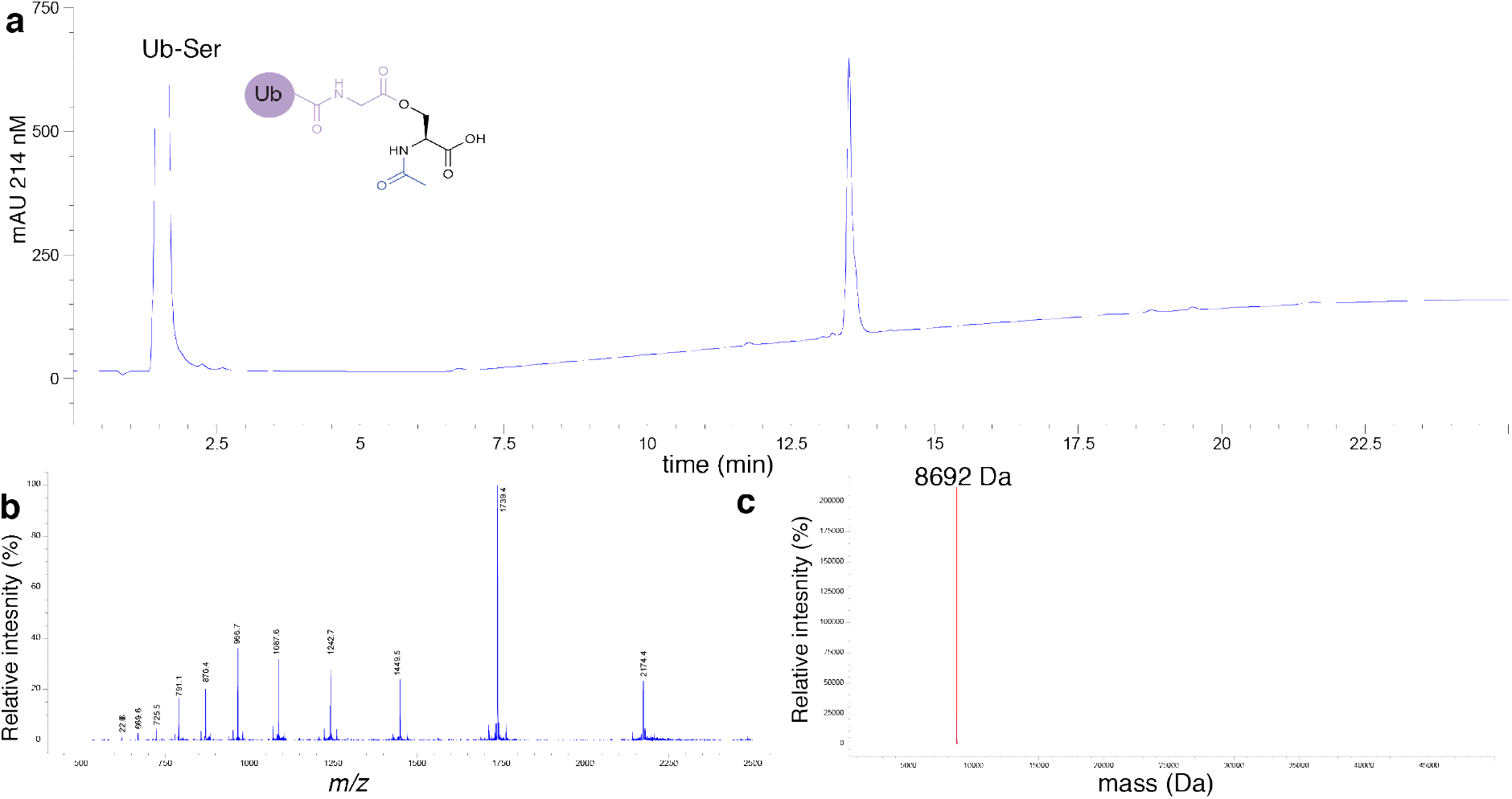
LC-MS characterization of purified Ub-Ser. **(a)** HPLC chromatogram for purified Ub-Ser monitoring at 214 nm. **(b)** Electrospray ionization mass spectrum for Ub-Ser. **(c)** Deconvoluted mass spectrum for Ub-Ser. Theoretical mass = 8694.13; observed mass = 8962 Da.

**Table S1.** Recombinant DUBS employed in this study. The known substrate corresponds to the substrate chosen in control experiments carried out to confirm DUB activity. If the DUB was expressed in truncated form then residue boundaries are stated (FL corresponds to full-length). DU number corresponds to MRC Reagents and Services internal reference number. Purity was approximated by Coomassie stained SDS-PAGE analysis. Concentration corresponds to that employed in MALDI-TOF assays unless otherwise stated.

| Enzyme | Known Substrate | Length | Tag | DU Number | Expression system | Purity | Average Mass | Concentration ng/μl | Concentration nM |
| --- | --- | --- | --- | --- | --- | --- | --- | --- | --- |
| USP1/UAF1 | K63 | FL | His | DU23056 | bacteria | 50% | 90502 | 18.0 | 198.9 |
| USP2 | K63 | FL | GST | DU13025 | bacteria | 50% | 72731 | 7.5 | 103.1 |
| USP4 | K48 | FL | His | DU14350 | bacteria | 40% | 111006 | 7.2 | 64.9 |
| USP5 | K63 | FL | His | DU15641 | bacteria | 90% | 98880 | 0.2 | 1.8 |
| USP6 | K63 | 529-1406 | GST | DU37745 | insect | 90% | 125292 | 0.2 | 1.4 |
| USP7 | K11 | FL | His | DU15644 | bacteria | 80% | 130714 | 0.1 | 0.7 |
| USP8 | K63 | FL | His | DU15645 | bacteria | 90% | 130789 | 180.0 | 1376.4 |
| USP9x | K11 | 1553-1995 | GST | DU20628 | bacteria | 50% | 78783 | 36.0 | 457.0 |
| USP10 | K48 | FL | His | DU15647 | bacteria | 80% | 91967 | 36.0 | 391.5 |
| USP11 | K48 | FL | GST | DU20016 | bacteria | 20% | 137621 | 12.0 | 87.2 |
| USP12 | K48 | FL | His | DU24373 | insect | 40% | 45127 | 12.0 | 265.9 |
| USP15 | K48 | FL | GST | DU19772 | bacteria | 90% | 137410 | 1.1 | 7.9 |
| USP16 | K63 | FL | GST | DU46239 | bacteria | 80% | 120394 | 3.6 | 29.9 |
| USP20 | K63 | FL | GST | DU12807 | bacteria | 40% | 128708 | 9.6 | 74.6 |
| USP21 | K48 | 196-565 | His | DU22384 | bacteria | 80% | 42852 | 7.2 | 168.0 |
| USP25 | K11 | FL | - | DU21610 | bacteria | 80% | 149037 | 1.8 | 12.1 |
| USP27x | K11 | FL | Dac | DU23206 | insect | 85% | 76453 | 36.0 | 470.9 |
| USP28 | K11 | FL | GST | DU20233 | bacteria | 60% | 149310 | 3.6 | 24.1 |
| USP30 | K48 | 57-517 | - | DU36294 | bacteria | 50% | 52995 | 72.0 | 1358.8 |
| USP36 | K11 | 81-461 | GST | DU4944 | bacteria | 80% | 69387 | 27.0 | 389.2 |
| USP45 | K48 | FL | GST | DU15681 | bacteria | 50% | 118554 | 60.0 | 506.1 |
| USP46 | K48 | FL | His | DU24347 | insect | 90% | 44711 | 30.0 | 671.0 |
| USP47 | K48 | FL | His | DU15682 | insect | 80% | 159545 | 3.0 | 18.8 |
| CYLD | K63 | FL | His | DU1834 | insect | 90% | 110553 | 72.0 | 651.3 |
| OTULIN | M1 | FL | - | DU43487 | bacteria | 95% | 40462 | 0.02 | 0.4 |
| OTU1 | None | FL | GST | DU36559 | bacteria | 90% | 65144 | 60.0 | 921.1 |
| OTUD1 | K63 | 270-481 | His | DU25080 | bacteria | 90% | 29718 | 0.2 | 6.1 |
| OTUD3 | K11 | FL | His | DU21336 | bacteria | 80% | 47566 | 3.6 | 75.7 |
| OTUD5 p177S | K63 | FL | - | DU21450 | bacteria | 70% | 84826 | 90.0 | 1061.1 |
| OTUD6A | None | FL | His | DU20889 | bacteria | 90% | 35743 | 60.0 | 1678.8 |
| OTUD6B | None | FL | GST | DU21140 | bacteria | 90% | 64148 | 60.0 | 935.4 |
| OTUB1 | K48 | FL | GST | DU19741 | bacteria | 90% | 58106 | 1.8 | 31.0 |
| OTUB2 | K63 | FL | GST | DU32795 | bacteria | 95% | 54035 | 0.4 | 6.7 |
| VCPIP1 | K48 | 25-561 | His | DU25040 | bacteria | 70% | 137252 | 36.0 | 262.3 |
| vOTU | K63 | 1 - 183 | GST | DU45351 | bacteria | >80% | 47660 | 0.02 | 0.4 |
| TRABID | K63 | 245-697 | His | DU22468 | bacteria | 50% | 54792 | 90.0 | 1642.7 |
| A20 | K48 | 1-366 | - | DU32912 | bacteria | 90% | 70843 | 9.0 | 127.1 |
| Cezanne | K11 | FL | GST | DU20899 | bacteria | 50% | 119719 | 0.04 | 0.3 |
| UCHL1 | Ub-W | FL | His | DU15693 | bacteria | 95% | 27267 | 0.3 | 11.0 |
| UCHL3 | Ub-W | FL | GST | DU21015 | bacteria | 80% | 53984 | 0.03 | 0.6 |
| UCHL5 | Ub-W | FL | GST | DU12810 | bacteria | 80% | 64300 | 210.0 | 3266.3 |
| BAP1 | Ub-W | FL | GST | DU12809 | bacteria | 70% | 107182 | 120.0 | 1119.7 |
| JOSD1 | None | FL | His | DU20956 | bacteria | 95% | 25641 | 60.0 | 2340.2 |
| JOSD2 | K48 | FL | His | DU20941 | bacteria | 90% | 23199 | 60.0 | 2586.6 |
| ATXN-3 | K63 | FL | GST | DU35448 | bacteria | 90% | 69225 | 60.0 | 866.8 |
| ATXN-3L | K63 | FL | His | DU23069 | bacteria | 70% | 43191 | 60.0 | 1389.3 |
| AMSH | K63 | 356-424 | - | DU44746 | bacteria | >80% | 74769 | 180.0 | 2407.7 |
| AMSH-LP | K63 | 265-436 | - | DU15780 | bacteria | 80% | 47266 | 0.1 | 1.9 |
| MINDY1 | K48 tetramer | 1 - 469 | MBP | DU59325 | bacteria | >80% | 96044 | 60.0 | 624.8 |
| MINDY2 | K48 | 241 - 504 | GST | DU55390 | bacteria | >80% | 58182 | 60.0 | 1031.3 |
| MINDY3 | K48 | FL | GST | DU47870 | bacteria | >85% | 76500 | 60.0 | 784.4 |
| MINDY4 | None | FL | GST | DU47893 | bacteria | >20% | 111100 | 60.0 | 540.1 |
| ZUFSP | K63 | 231-578 | GST | DU53621 | bacteria | 90% | 66600 | 60.0 | 901.0 |
| UAF1 | None | FL | His | DU39055 | insect | 80% | 81023 | 3.0 | 37.0 |

## References

1 Oh, E., Akopian, D. & Rape, M. Principles of Ubiquitin-Dependent Signaling. Annu Rev Cell Dev Biol 34, 137–162, doi:10.1146/annurev-cellbio-100617-062802 (2018).

2 Hershko, A. & Ciechanover, A. The ubiquitin system. Annu Rev Biochem 67, 425–479, doi:10.1146/annurev.biochem.67.1.425 (1998).

3 Deshaies, R. J. & Joazeiro, C. A. RING domain E3 ubiquitin ligases. Annu Rev Biochem 78, 399–434, doi:10.1146/annurev.biochem.78.101807.093809 (2009).

4 Zheng, N. & Shabek, N. Ubiquitin Ligases: Structure, Function, and Regulation. Annu Rev Biochem 86, 129–157, doi:10.1146/annurev-biochem-060815-014922 (2017).

5 Pao, K. C. et al. Activity-based E3 ligase profiling uncovers an E3 ligase with esterification activity. Nature 556, 381–385, doi:10.1038/s41586-018-0026-1 (2018).

6 Kwon, Y. T. & Ciechanover, A. The Ubiquitin Code in the Ubiquitin-Proteasome System and Autophagy. Trends Biochem Sci 42, 873–886, doi:10.1016/j.tibs.2017.09.002 (2017).

7 Clague, M. J., Urbe, S. & Komander, D. Breaking the chains: deubiquitylating enzyme specificity begets function. Nat Rev Mol Cell Biol 20, 338–352, doi:10.1038/s41580-019-0099-1 (2019).

8 Goto, E. et al. c-MIR, a human E3 ubiquitin ligase, is a functional homolog of herpesvirus proteins MIR1 and MIR2 and has similar activity. J Biol Chem 278, 14657–14668, doi:10.1074/jbc.M211285200 (2003).

9 Cadwell, K. & Coscoy, L. Ubiquitination on nonlysine residues by a viral E3 ubiquitin ligase. Science 309, 127–130, doi:10.1126/science.1110340 (2005).

10 Wang, X. et al. Ubiquitination of serine, threonine, or lysine residues on the cytoplasmic tail can induce ERAD of MHC-I by viral E3 ligase mK3. J Cell Biol 177, 613–624, doi:10.1083/jcb.200611063 (2007).

11 Jin, L., Williamson, A., Banerjee, S., Philipp, I. & Rape, M. Mechanism of ubiquitin- chain formation by the human anaphase-promoting complex. Cell 133, 653–665, doi:10.1016/j.cell.2008.04.012 (2008).

12 Wang, X. et al. Ube2j2 ubiquitinates hydroxylated amino acids on ER-associated degradation substrates. J Cell Biol 187, 655–668, doi:10.1083/jcb.200908036 (2009).

13 Grill, B., Murphey, R. K. & Borgen, M. A. The PHR proteins: intracellular signaling hubs in neuronal development and axon degeneration. Neural Dev 11, 8, doi:10.1186/s13064-016-0063-0 (2016).

14 Tokunaga, F. et al. Involvement of linear polyubiquitylation of NEMO in NF-kappaB activation. Nat Cell Biol 11, 123–132, doi:10.1038/ncb1821 (2009).

15 Kelsall, I. R., Zhang, J., Knebel, A., Arthur, J. S. C. & Cohen, P. The E3 ligase HOIL-1 catalyses ester bond formation between ubiquitin and components of the Myddosome in mammalian cells. Proc Natl Acad Sci U S A 116, 13293–13298, doi:10.1073/pnas.1905873116 (2019).

16 Sun, H., Meledin, R., Mali, S. M. & Brik, A. Total chemical synthesis of ester-linked ubiquitinated proteins unravels their behavior with deubiquitinases. Chem Sci 9, 1661–1665, doi:10.1039/c7sc04518b (2018).

17 Ritorto, M. S. et al. Screening of DUB activity and specificity by MALDI-TOF mass spectrometry. Nat Commun 5, 4763, doi:10.1038/ncomms5763 (2014).

18 Virdee, S., Ye, Y., Nguyen, D. P., Komander, D. & Chin, J. W. Engineered diubiquitin synthesis reveals Lys29-isopeptide specificity of an OTU deubiquitinase. Nat Chem Biol 6, 750–757, doi:10.1038/nchembio.426 (2010).

19 Tam, J. P., Yu, Q. & Lu, Y. A. Tandem peptide ligation for synthetic and natural biologicals. Biologicals 29, 189–196, doi:10.1006/biol.2001.0292 (2001).

20 Hadari, T., Warms, J. V., Rose, I. A. & Hershko, A. A ubiquitin C-terminal isopeptidase that acts on polyubiquitin chains. Role in protein degradation. J Biol Chem 267, 719–727 (1992).

21 Akutsu, M., Ye, Y., Virdee, S., Chin, J. W. & Komander, D. Molecular basis for ubiquitin and ISG15 cross-reactivity in viral ovarian tumor domains. Proc Natl Acad Sci U S A 108, 2228–2233, doi:10.1073/pnas.1015287108 (2011).

22 Tran, H., Hamada, F., Schwarz-Romond, T. & Bienz, M. Trabid, a new positive regulator of Wnt-induced transcription with preference for binding and cleaving K63- linked ubiquitin chains. Genes Dev 22, 528–542, doi:10.1101/gad.463208 (2008).

23 Jin, J. et al. Epigenetic regulation of the expression of Il12 and Il23 and autoimmune inflammation by the deubiquitinase Trabid. Nat Immunol 17, 259–268, doi:10.1038/ni.3347 (2016).

24 Licchesi, J. D. et al. An ankyrin-repeat ubiquitin-binding domain determines TRABID’s specificity for atypical ubiquitin chains. Nat Struct Mol Biol 19, 62–71, doi:10.1038/nsmb.2169 (2011).

25 Zhu, Y. et al. Trabid inhibits hepatocellular carcinoma growth and metastasis by cleaving RNF8-induced K63 ubiquitination of Twist1. Cell Death Differ 26, 306–320, doi:10.1038/s41418-018-0119-2 (2019).

26 Mevissen, T. E. T. & Komander, D. Mechanisms of Deubiquitinase Specificity and Regulation. Annu Rev Biochem 86, 159–192, doi:10.1146/annurev-biochem-061516-044916 (2017).

27 Kristariyanto, Y. A., Abdul Rehman, S. A., Weidlich, S., Knebel, A. & Kulathu, Y. A single MIU motif of MINDY-1 recognizes K48-linked polyubiquitin chains. EMBO Rep 18, 392–402, doi:10.15252/embr.201643205 (2017).

28 Kwasna, D. et al. Discovery and Characterization of ZUFSP/ZUP1, a Distinct Deubiquitinase Class Important for Genome Stability. Mol Cell 70, 150–164 e156, doi:10.1016/j.molcel.2018.02.023 (2018).

29 Hermanns, T. et al. A family of unconventional deubiquitinases with modular chain specificity determinants. Nat Commun 9, 799, doi:10.1038/s41467-018-03148-5 (2018).

30 Haahr, P. et al. ZUFSP Deubiquitylates K63-Linked Polyubiquitin Chains to Promote Genome Stability. Mol Cell 70, 165–174 e166, doi:10.1016/j.molcel.2018.02.024 (2018).

31 Hewings, D. S. et al. Reactive-site-centric chemoproteomics identifies a distinct class of deubiquitinase enzymes. Nat Commun 9, 1162, doi:10.1038/s41467-018-03511-6 (2018).

32 Sato, Y. et al. Structural basis for specific cleavage of Lys 63-linked polyubiquitin chains. Nature 455, 358–362, doi:10.1038/nature07254 (2008).

33 Geurink, P. P., El Oualid, F., Jonker, A., Hameed, D. S. & Ovaa, H. A general chemical ligation approach towards isopeptide-linked ubiquitin and ubiquitin-like assay reagents. Chembiochem 13, 293–297, doi:10.1002/cbic.201100706 (2012).

34 Plechanovova, A. et al. Mechanism of ubiquitylation by dimeric RING ligase RNF4. Nat Struct Mol Biol 18, 1052–1059, doi:10.1038/nsmb.2108 (2011).

35 Nijman, S. M. et al. A genomic and functional inventory of deubiquitinating enzymes. Cell 123, 773–786, doi:10.1016/j.cell.2005.11.007 (2005).

36 Burnett, B., Li, F. & Pittman, R. N. The polyglutamine neurodegenerative protein ataxin-3 binds polyubiquitylated proteins and has ubiquitin protease activity. Hum Mol Genet 12, 3195–3205, doi:10.1093/hmg/ddg344 (2003).

37 Matos, C. A., de Macedo-Ribeiro, S. & Carvalho, A. L. Polyglutamine diseases: the special case of ataxin-3 and Machado-Joseph disease. Prog Neurobiol 95, 26–48, doi:10.1016/j.pneurobio.2011.06.007 (2011).

38 Masino, L. et al. Domain architecture of the polyglutamine protein ataxin-3: a globular domain followed by a flexible tail. FEBS Lett 549, 21–25, doi:10.1016/s0014-5793(03)00748-8 (2003).

39 Seki, T. et al. JosD1, a membrane-targeted deubiquitinating enzyme, is activated by ubiquitination and regulates membrane dynamics, cell motility, and endocytosis. J Biol Chem 288, 17145–17155, doi:10.1074/jbc.M113.463406 (2013).

40 Wu, X. et al. JOSD1 inhibits mitochondrial apoptotic signalling to drive acquired chemoresistance in gynaecological cancer by stabilizing MCL1. Cell Death Differ, doi:10.1038/s41418-019-0339-0 (2019).

41 Wang, X. et al. JOSD1 Negatively Regulates Type-I Interferon Antiviral Activity by Deubiquitinating and Stabilizing SOCS1. Viral Immunol 30, 342–349, doi:10.1089/vim.2017.0015 (2017).

42 Weeks, S. D., Grasty, K. C., Hernandez-Cuebas, L. & Loll, P. J. Crystal structure of a Josephin-ubiquitin complex: evolutionary restraints on ataxin-3 deubiquitinating activity. J Biol Chem 286, 4555–4565, doi:10.1074/jbc.M110.177360 (2011).

43 Renatus, M. et al. Structural basis of ubiquitin recognition by the deubiquitinating protease USP2. Structure 14, 1293–1302, doi:10.1016/j.str.2006.06.012 (2006).

44 Emmerich, C. H. et al. Activation of the canonical IKK complex by K63/M1-linked hybrid ubiquitin chains. Proc Natl Acad Sci U S A 110, 15247–15252, doi:10.1073/pnas.1314715110 (2013).

45 Geurink, P. P. et al. Development of Diubiquitin-Based FRET Probes To Quantify Ubiquitin Linkage Specificity of Deubiquitinating Enzymes. Chembiochem 17, 816–820, doi:10.1002/cbic.201600017 (2016).

46 Kelley, L. A., Mezulis, S., Yates, C. M., Wass, M. N. & Sternberg, M. J. The Phyre2 web portal for protein modeling, prediction and analysis. Nat Protoc 10, 845–858, doi:10.1038/nprot.2015.053 (2015).

47 Brownell, J. E. et al. Substrate-assisted inhibition of ubiquitin-like protein-activating enzymes: the NEDD8 E1 inhibitor MLN4924 forms a NEDD8-AMP mimetic in situ. Mol Cell 37, 102–111, doi:10.1016/j.molcel.2009.12.024 (2010).

48 De Cesare, V. et al. The MALDI-TOF E2/E3 Ligase Assay as Universal Tool for Drug Discovery in the Ubiquitin Pathway. Cell Chem Biol 25, 1117–1127 e1114, doi:10.1016/j.chembiol.2018.06.004 (2018).

